# The cerebellum contributes to tonic-clonic seizures by altering neuronal activity in the ventral posteromedial nucleus (VPM) of the thalamus

**DOI:** 10.1101/2021.10.03.462953

**Authors:** Jaclyn Beckinghausen, Joshua Ortiz-Guzman, Tao Lin, Benjamin Bachman, Yu Liu, Detlef H. Heck, Benjamin R. Arenkiel, Roy V. Sillitoe

## Abstract

Thalamo-cortical networks are central to seizures, yet it’s unclear how these circuits initiate the seizures. Here, we test the hypothesis that a facial region of the thalamus, the VPM, is a source of convulsive, tonic-clonic seizures. We devised an *in vivo* optogenetic mouse model to elicit tonic-clonic seizures by driving convergent input to the VPM. With viral tracing, we show dense cerebellar and cerebral cortical afferent input to the VPM. Lidocaine microinfusions into the cerebellar nuclei selectively block seizure initiation. We perform single-unit electrophysiology recordings during awake, convulsive seizures to define the local activity of thalamic neurons before, during, and after seizure onset. We find highly dynamic activity with biphasic properties, raising the possibility that heterogenous activity patterns promote seizures. These data reveal the VPM as a source of tonic-clonic seizures, with cerebellar input providing the predominant signals.

## Introduction

At their worst, seizures can be fatal, and at best, they significantly decrease a patient’s quality of life. Regardless of whether a traumatic brain injury or a specific genetic mutation is responsible for starting the seizure, about a third of patients experience persistent seizures despite having pharmacological, dietary, or surgical interventions^1–9^. In over 60% of cases, the catalyst for initial seizure precipitation is unclear and the causes of such “cryptogenic” seizures remain unknown^10^. As a consequence, it is difficult to optimize therapeutic options for seizure patients^3^. Thus, there is a pressing need to uncover how abnormal seizure activity arises and how it impacts behavior.

Of all the types of seizures, convulsive tonic-clonic seizures (TC) are perhaps the most disruptive and generally feared among patients and caregivers. Unfortunately, these are the most common class of seizures in disorders collectively called epilepsy^11^. Two brain regions that have received the majority of focus in these studies are the cerebral cortex and thalamus, as these reciprocally connected regions form a hypersynchronous, self-perpetuating loop in many forms of epilepsy. Numerous studies have demonstrated that the cortico-thalamic circuit is important in seizures; specifically, such studies point to the cerebral cortex as a main site of initiation^12–14^. However, with a large portion of epilepsy patients being refractory to treatment, stemming from attempts to target these well-studied circuits, it may be critical to narrow in on specific seizure foci and to expand the seizure circuit to determine whether other combinations of neural networks contribute to ictogenesis. We therefore set out to delineate specific focal areas within the cortico-thalamic loop and to identify extra-thalamic inputs that contribute to seizure initiation, and subsequently interrogated the neural activity that may underlie convulsive seizures in awake, seizing animals.

Deeper examination of areas outside the cerebral cortex revealed that seizures can arise from a number of subcortical regions, including the amygdala, hippocampus, and basal ganglia^15–18^. Other regions, such as the cerebellum, are thought to play some role in seizures, but the extent of their contribution is controversial^19^. The majority of these regions project to the thalamus, which is not only a well-established area of information integration, but is also understood to serve as a key gateway for mediating seizures that depend upon these various regions^20–22^. Yet, the precise role of individual thalamic nuclei and their unique combination of inputs on the generation of abnormal brain activity remains unclear. Pharmacological inactivation studies of thalamic targets, namely the motor or sensory cortices, demonstrate a clear requirement for somatosensory areas in both initiation and maintenance of spike-wave discharges (SWD), a hallmark EEG pattern of absence seizures^12, 23–27^. Similarly, the pattern of activation based on human MRI and analysis of multi-electrode recordings in genetic rat models identify SWDs and activity changes specifically in the peri-oral region of the somatosensory cortex prior to spread in other regions of the cortico-thalamic loop^12, 23, 28^. While such findings have been primarily studied in absence seizures, the possibility of an epileptic origin in peri-oral circuits is particularly intriguing for TC seizures, as facial clonus is one of the first behavioral readouts observed in this dynamic phenotype. We therefore hypothesize that specific cortico-thalamic loops involving facial regions may be particularly important in seizure generation. Furthermore, in patients with convulsive seizures, recent human studies reported thalamic changes before cortical involvement based on electrode recordings in putative deep brain stimulation patients^29^. These findings thus suggest that the thalamus leads the cortex in generalized seizures, whereas frontal seizures exhibit activity in the opposite direction^29^. With particular interest in cryptogenic, generalized seizures, we therefore chose to test circuits in thalamic regions that are known to be tightly interconnected with the facial somatosensory cortex.

The ventral posteromedial thalamic nucleus (VPM) projects directly to layers 4, 5B, and 6A of the primary facial somatosensory cortex and receives primarily trigeminal input from the head, face, and oral regions^30–33^. Known to be intimately involved in the production of spindle waves, a 7-15Hz rhythm occurring during sleep and seizures^34^, the VPM has been shown to be an effective locus in aborting post-stroke epileptic seizures^35^. Due to its reciprocal connections with the reticular thalamic nucleus as well as direct innervation of both excitatory and inhibitory neurons in the facial somatosensory cortex, the VPM is a strong candidate for initiating generalized seizure activity. Using optogenetics to mimic cryptogenic seizures, we test whether activation of afferent inputs specifically to the VPM, but not to other surrounding thalamic nuclei, drives severe tonic-clonic seizures. We also used viral tracing to identify the specific neural projections that may contribute to ictogenesis: specifically, those projections that are monosynaptically connected to the VPM.

The cerebellum is one such source of direct projections to the VPM. Historically, a cerebellar role in seizures has been controversial. Earlier observations of seizures, especially hemi-facial spasms (HFS), in patients with cerebellar gangliomas^36, 37^ inspired the use of the cerebellar cortex as a deep brain stimulation (DBS) target for epilepsy patients dating back to the 1970’s^38, 39^. Despite its success in alleviating seizures in early clinical tests, ethical dilemmas in these studies and variability in success with subsequent trials due to the previously unknown complexity of the cerebellar circuitry led to shift in attention away from the cerebellum as a seizure locus. Still, cerebellar pathology is consistently observed in patients with numerous disorders containing a common denominator: seizures. For example, in Unverrict Lundborg Disease and familial cortical myoclonic tremor and epilepsy, both of which exhibit seizures, degeneration in cerebellar granule cells, Purkinje cells, and overall cerebellar atrophy are observed in human patients^40, 41^. Diffusion tensor imaging has shown abnormal white matter connectivity within the cerebellum in idiopathic epilepsy^42^. With the growing evidence of cerebellar damage in human patients with seizures^43–46^, plus its involvement in the induction^47, 48^ and resolving of seizures^43, 49–53^ when different stimulation paradigms, this previously disregarded seizure-related brain region may unlock new therapeutic potential. Although recent studies have demonstrated behavioral and electrophysiological control of seizures using the cerebellum in genetic mouse models of absence epilepsy^50, 53, 54^, we have a limited understanding about how to (1) elicit a severe, TC seizure without using drugs, chemicals, current, or gene-specific mutations, or (2) how cerebellar circuitry and function fit into a broader seizure brain network. With our optogenetics model, we test the role of the VPM in seizures and address whether the cerebellum may contribute to the initiation of TC seizures in this mouse model.

## Results

### *Ntsr1^Cre^* transgenic mice drive reporter expression in brain regions involved in seizures

*Ntsr1* encodes for a protein that is expressed primarily in the gastrointestinal tract, but is also widely distributed throughout the brain^55^. A previously generated transgenic mouse model^56, 57^ demonstrated that the gene’s regulatory sequences can be used to drive *Cre* expression across different brain regions including layers 5 and 6 of the cerebral cortex, the thalamus, and the ventral tegmental area^58, 59^. Because a key aspect of our study relies on the identities of *Cre* expressing neurons that may contribute to ictogenesis, we first set out to both confirm and expand upon previous characterizations of this line. To evaluate the *Ntsr1* transgene expression across the whole brain, we crossed *Ntsr1^Cre^* male mice to two different reporter lines: Sun1 (*Ntsr1^Cre^;ROSA^lox-stop-lox-Sun^*^1^), a small nuclear envelope protein expressed in the nuclear membrane (Fig 1a), and TdTomato (*Ntsr1^Cre^;ROSA^lox-stop-lox-TdTomato^*), which typically has the advantage of brightly labeling cells and showing their architecture in full due to the intensity of its fluorescent emission (Fig 1f). The concomitant use of these lines allowed us to distinguish between areas containing *Cre* recombined resident neurons (Sun1 positive) versus areas that receive input from *Ntsr1^Cre^*-positive neurons and therefore exhibit fluorescence specifically from their axonal processes and terminals (TdTomato positive).

**Figure 1:**
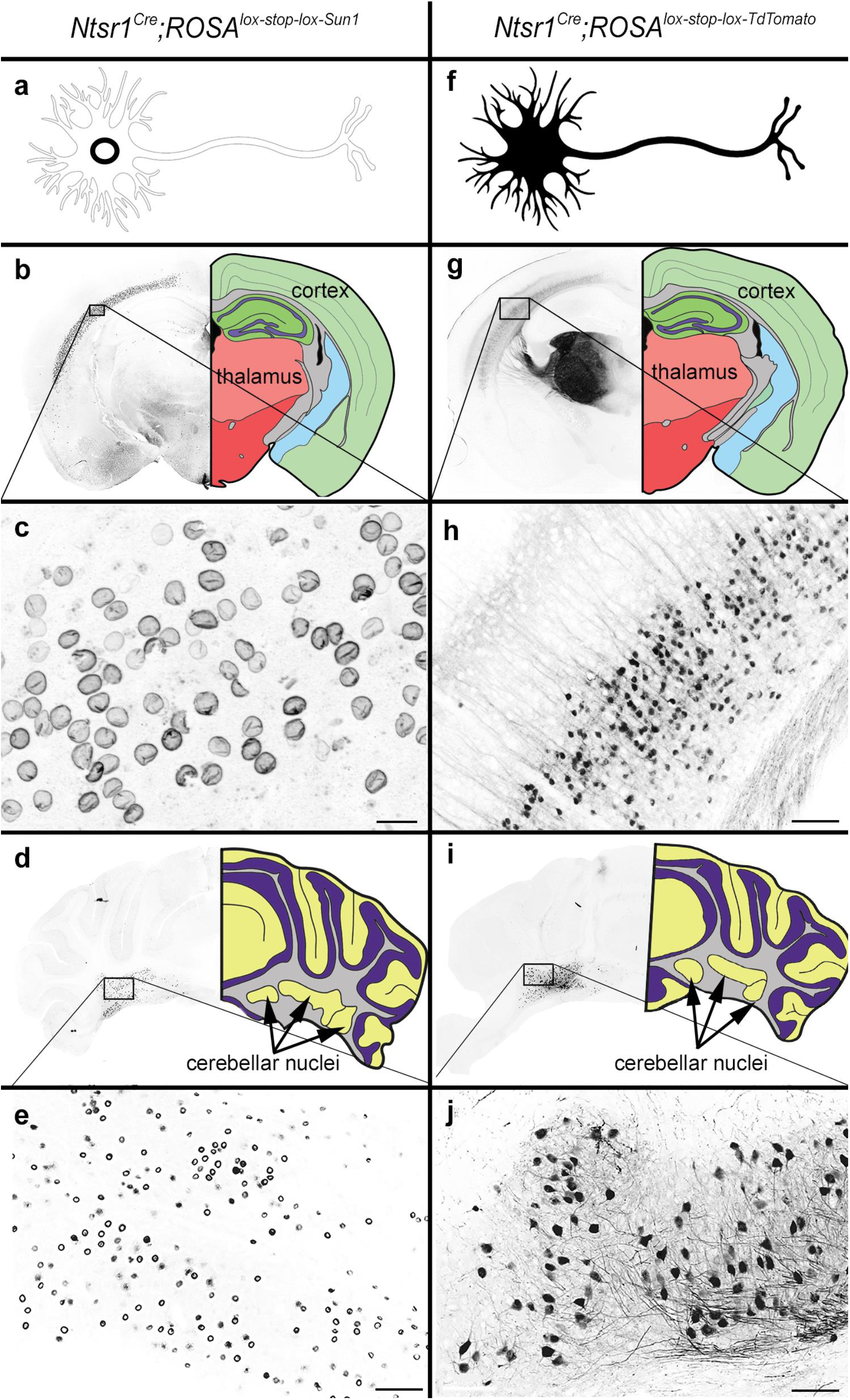
Ntsr1^Cre^ is expressed in corticothalamic layer 6, cerebellar nuclei, and the axons and fibers of passage within the thalamic nuclei. (a) Schematic of Sun1 expression, which labels the nuclear envelope of Cre-expressing neurons. (b) Expression of Sun1 is seen in layer 6 of the cerebral cortex (black box, high power image in (c)), but not in thalamic nuclei. (c) High power image of cortical neurons expressing nuclear Sun1. Scale bar = 20um. (d) Cerebellar nuclei neurons in the dentate and interposed nuclei express Sun1. (e) High-power image of Sun1 in the cerebellar nuclei, showing the specificity of fluorescence for the nuclear membrane. Scale bar = 100um. (f) Schematic of Tdtomato expression, which fills the entire cell including dendrites and axonal processes. (g) Expression of Tdtomato is observed in layer 6 of the cerebral cortex (black box, high power image in (h)) as well as in the thalamus. However, in context with the lack of Sun1 expression in (b), this thalamic TdTomato signal can be identified as input from other Cre-expressing areas. (h) High power image demonstrating TdTomato expression in cortical layer 5/6 neurons. Scale bar = 100um. (i) Cerebellar nuclei neurons in the dentate and interposed nuclei express TdTomato. (j) High power magnification of cerebellar nuclei neurons with Tdtomato fluorescence in the somas as well as in axons exiting the cerebellum. Scale bar = 100um.

Coronal sections from *Ntsr1^Cre^;ROSA^lox-stop-lox-Sun^*^1^ mouse brains showed Sun1 nuclear membrane labeling in layer 5/6 of the cerebral cortex (Fig 1b, box; Fig 1c). This particular region with *Ntsr1^Cre^*-positive expression was studied previously and those neurons were identified as excitatory, corticothalamic projection neurons^60^. Another area that was positive for Sun1 reporter expression was the cerebellar nuclei (Fig 1d,e), which was again previously reported to be a region in which *Ntsr1^Cre^* drives expression in excitatory, VGLUT2 (vesicular-glutamate transporter 2)-expressing projection neurons^61^. However, we observed that the expression within the cerebellar nuclei was not evenly distributed: the medial cerebellar nuclei (fastigial nucleus) contained very few *Cre*-positive neurons. In contrast, we found that the reporter signal was heavily localized to projection neurons in the interposed (central) and dentate (lateral) nuclei. This is an important consideration when manipulating cerebellar output, as each nucleus has distinct downstream targets^62–64^. Although the cerebral cortex and cerebellum contained the most prominent distribution of cells with reporter expression, we also observed Sun1 labeling in the VTA^58^, caudoputamen and superior colliculus, as well as sparse labeling in hippocampal and thalamic neurons, and in a small number of cells that were scattered throughout the pontine nuclei and various brainstem nuclei (Fig S1ai-av).

By crossing the *Ntsr1^Cre^* mouse to a TdTomato reporter line, we were able to mark and clearly label somata as well as axonal compartments in each of the regions also marked by Sun1. This allowed us to distinguish between regions in which *Cre-*expressing neurons originate (that is, where their somas are located) versus regions that these neurons project to. In addition, labeling of the neurons in their entirety is important for colocalization analyses. To better characterize the identity of these neurons, we therefore stained 40um sections from our TdTomato reporter samples with NeuN, a pan-neuronal marker, NFH, an axonal and synaptic marker of neurons, and calbindin, a calcium binding protein that labels unique populations of neurons in different regions of the brain. In the cerebral cortex, calbindin is expressed in non-pyramidal cells^65^. Thalamic neurons with calbindin immunoreactivity identify subpopulations within functionally distinct nuclei^66^, whereas in the cerebellum, calbindin is highly expressed in GABAergic Purkinje cells that project to the cerebellar nuclei neurons. In the *Ntsr1^Cre^* cross with the TdTomato line, TdTomato positive cells colocalized with NeuN but not calbindin in the cortex (Fig S2a-f). These data support previous observations showing that *Ntsr1^Cre^* drives recombination of a reporter allele in excitatory pyramidal cells^67^. We also co-stained with tyrosine hydroxylase, a dopaminergic cell marker, to confirm the identity of the VTA-localized, *Ntsr1^Cre^* marked neurons as dopaminergic (Fig S2g-i). In the cerebellum, we confirmed previous reports of *Ntsr1^Cre^*-positive neurons as excitatory and NFH-positive cells that receive input from calbindin-positive, inhibitory Purkinje cells (Fig S3a-o). We also observed a variable number of *Ntsr1^Cre^-*positive Purkinje cells (Fig S3p-r) that were distributed throughout the structure without an obvious pattern of localization. In any given 40um section through cerebellar cortex, we could identify anywhere from 0 to 20 labeled Purkinje cells scattered throughout the section. Importantly, the low number of marked Purkinje cells was unlikely to influence our strategy for modulating neuronal activity with *Ntsr1^Cre^* (see below). We also found that *Ntsr1^Cre^* marks a subset of molecular layer interneurons in lobules IX and X (Fig S3s-u).

While the TdTomato cross confirmed the presence of *Ntsr1^Cre^* recombination in cortico-thalamic (Fig 1g-h, Fig S1 bi-bvi, Fig S2a-f) and cerebellar nuclei projection neurons (Fig 1i-j, Fig S1c), the most notable difference between our Sun1 and TdTomato reporter mouse lines was the pattern of staining in thalamic sub-nuclei: while the thalamus was specifically devoid of fluorescence in the Sun1 mouse line (Fig 1b, Fig S1ai-aiii), the fluorescence present in thalamic nuclei of the TdTomato mouse was intense (Fig 1g, Fig S1bi-bvi). These data together suggest that the resident thalamic neurons themselves do not contain *Ntsr1^Cre^* expression. In fact, high power magnification images of the thalamus in the TdTomato mice reveals “pockets” where the fluorescent signal is missing (Fig 2b). We predicted that these ovoid profiles without staining were likely the somata of resident neurons. To further investigate these “signal void” pockets, we examined high-power images of calbindin and NeuN immunoreactivity. As hypothesized, th absence of the TdTomato signal corresponded with both calbindin and NeuN labeling in the cell body (Fig S4). Therefore, we conclude that the reporter expression observed in the thalamus of the *Ntsr1^Cre^;ROSA^lsl-TdTomato^* mice is the result of marking the incoming axons and fibers of passage that originate in other areas where *Ntsr1^Cre^* labels the somata. An example of such fibers providing the robust thalamic reporter signal can be seen in supplemental Figure 1c, which shows a dense bundle of cerebellar nuclei axons exiting the cerebellum to innervate the various thalamic nuclei as well as other brain regions.

**Figure 2:**
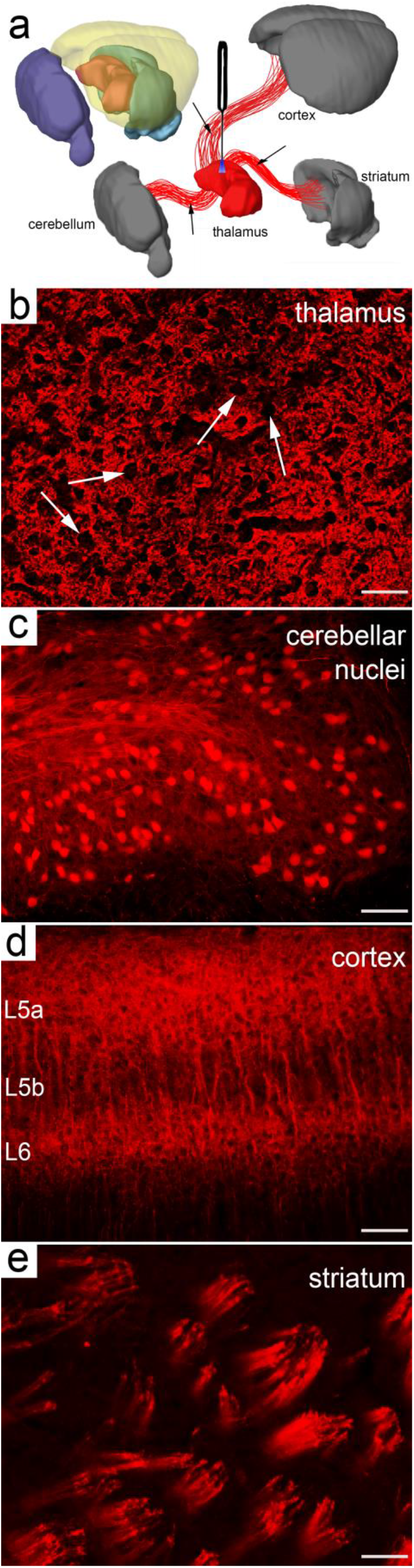
Optogenetic modulation of thalamus instead targets input from extra-thalamic regions. (a) Because the local thalamic neurons themselves do not express *Cre* in *Ntsr1^Cre^* animals, manipulation of their activity is indirect by modulating *Cre*-expressing axons and fibers of passage from regions outside the thalamus itself. Images of brain regions adapted and modified from BrainExplorer2, Allen Brain Atlas. (b-e) High power images of *Cre*-induced reporter expression patterns in the thalamus (scale bar = 50um) (b), cerebellar nuclei (scale bar = 100um) (c), cortex (scale bar = 100um) (d), and striatum (scale bar = 50um) (e). Note the distinction of the expression pattern of the thalamus compared to other regions (b), where fibers but not the resident neurons have expression and these areas are depicted by absence of localized fluorescence (arrows).

To gain optogenetic control over these genetically marked neurons, we next crossed the *Ntsr1^Cre^* mice to the *ROSA^lsl-Channelrhodopsin^* (hereafter referred to simply as ChR2) reporter strain. The offspring of this cross carried light-sensitive neuronal cell bodies in the aforementioned *Cre*-expressing brain regions as well as in the axons projecting to, or passing through, the thalamic nuclei. Based on this expression profile, we were able to deliver 465nm light to the thalamic nuclei in order to optogenetically modulate thalamic neuron activity indirectly by activating the axons and terminals of their presynaptic, ChR2-expressing afferent inputs (Fig 2a).

### High-frequency light pulses to the VPM elicit tonic-clonic seizures in *Ntsr1^Cre^;ChR2* mice

The thalamus is a critical structure in the seizure network^20–22^, although the contribution of its different nuclei to mediating seizures is not fully understood. Using the *Ntsr1^Cre^;ChR2* mice, we targeted blue light to manipulate the thalamic nuclei, as each of its major divisions were densely innervated by a convergence of axons from multiple *Cre*-expressing brain regions. We found that unilateral delivery of 30Hz pulses of light to the ventral posteromedial nucleus (VPM) elicits a robust, TC seizure phenotype (n=24 mice, supp video 1,2). Bilateral stimulation or co-stimulation with other *Cre-*expression regions, such as the cerebellar nuclei, did not enhance or reduce the severity of this phenotype (n=4 mice, supp video 3). Interestingly, stimulating the surrounding nuclei with the same parameters, including the posterior thalamic nucleus (PO, n=3 mice), the ventroposterior lateral nucleus (VPL, n=3 mice), the reticular thalamic nucleus (RTN, n=3 mice), and the lateral dorsal nucleus (LD, n=2 mice) did not elicit overt behavioral changes (supp video 4). We quantified the relative fluorescence in each of these individual thalamic nuclei (n=4 mice, 5-10 slices per mouse) to determine whether the non-responsive nuclei received less ChR2-expressing input. While some nuclei, such as the RTN, PO, and CL, had significantly less fluorescence (Dunnett’s multiple comparisons test, Table 1), which may account for the absence of behavior upon their stimulation, other nuclei like the VPL contained comparable levels of ChR2 input as the VPM (Fig 3a,b, Dunnett’s multiple comparisons test, Table 1). These data suggest that the initiation of seizures in *Ntsr1^Cre^;ChR2* mice requires the modulation of unique sets of inputs that converge within the VPM and argues against a global activation of general input pathways.

**Figure 3:**
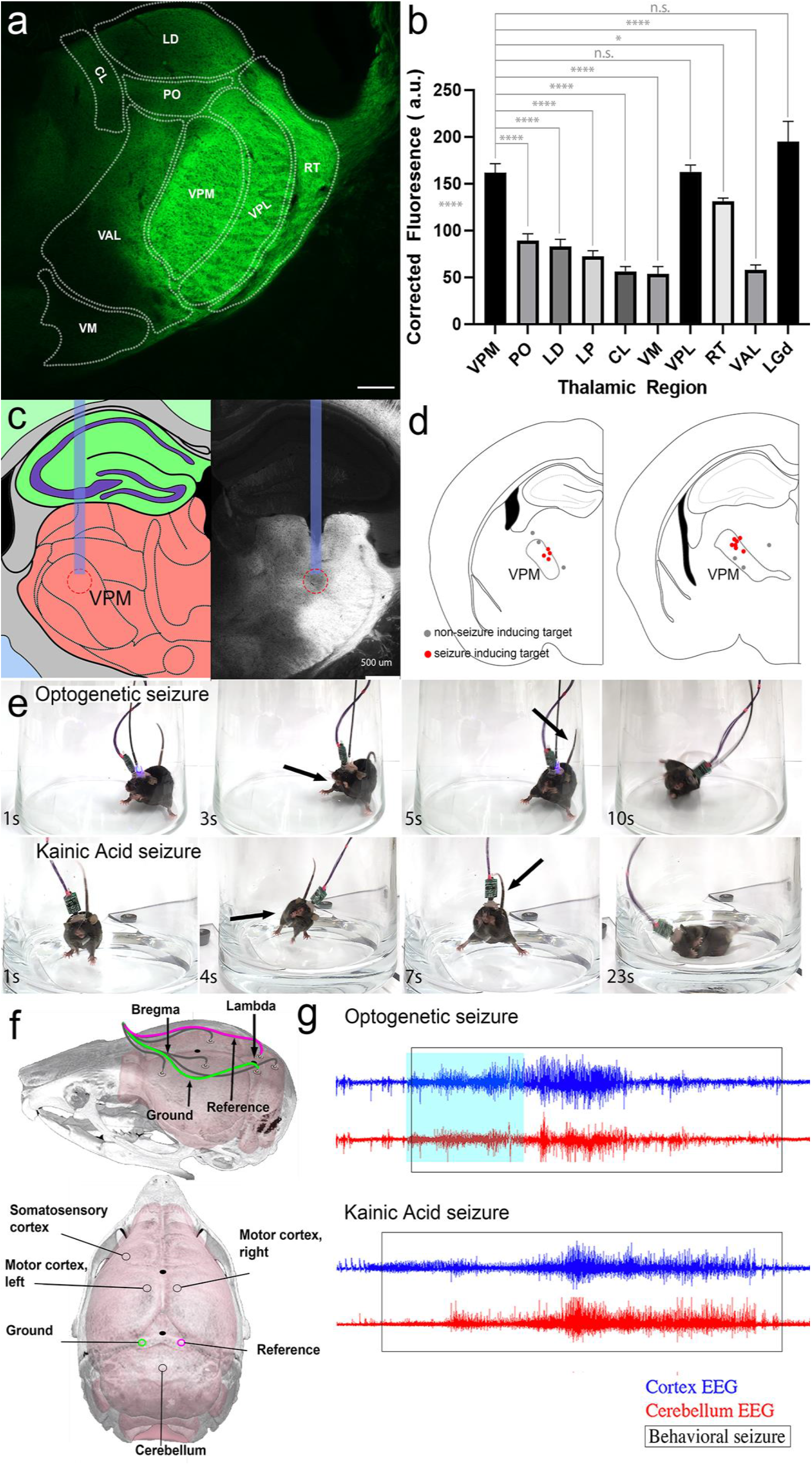
Seizure-inducing stimulations are specific to the VPM. (a) Representative segmentation of thalamic nuclei (coronal) showing relative fluorescence of various nuclei in *Ntsr1^Cre^;ROSA^lox-stop-lox-ChR^*^2^*^-YFP^* mice. (b) Quantification of fluorescent intensity of various thalamic nuclei, some of which are segmented in (a), demonstrates that the specificity of VPM as a seizure locus is not due to biased *Cre* expression, as no behavior is elicited even when VPL is targeted. A Dunnett’s multiple comparison’s test determined significance between various thalamic nuclei compared to the VPM as follows: VPM vs PO (p<0.0001), VPM vs LD (p<0.0001), VPM vs LP (p<0.0001), VPM vs CL (p<0.0001), VPM vs VM (p<0.0001), VPM vs VPL (ns), VPM vs RT (p=0.0156), VPM vs VAL (p<0.0001), VPM vs LGd (ns). (c) Example of post-hoc targeting of fiber optics to the VPM as demonstrated both by the electrode track and an area of photobleaching of the ChR2-YFP signal. (d) Targeting of the fiber optics are examined in mice at the end of experimentation. Each red dot represents the end of a fiber in a single mouse that successfully exhibited stage 6 seizures. Grey dots depict targeting that did not elicit seizures. (e) Example of the root mean square of left cortical EEG in a single mouse during multiple seizure stimulation periods (blue overlays, 30s stimulations). Each subsequent stimulation period begins 2 minutes following the termination of the previous stimulation. Behavioral seizures are marked by the yellow rectangles, which coincide with high amplitude EEG activity. Here, we see that stimulations immediately following a previously elicited seizure (within 2 minutes) do not induce subsequent seizures. Instead, mice experience a refractory period in which additional delivery of light does not result in behavioral abnormalities. Scale bars mark two instances in which the mouse only experiences a seizure 4 minutes and 6 minutes following a previous seizure, but not 2 minutes after a seizure.

*Post-hoc* anatomical examination of fiber optic targeting further identified a specific locus within the VPM bordering the PO that reliably elicited seizures (Fig 3c,d, n=7 mice). Targeting of fibers to other regions within the VPM did not elicit a behavioral change. These data indicate that a discrete set of VPM inputs are required for seizure initiation. With repeated stimulation, we determined that all *Ntsr1^Cre^;ChR2* mice with accurate VPM fiber targeting eventually exhibited stage 6 seizures according to a modified Racine Scale^68^ (Table 2), a widely used template for categorization and ranking of rodent seizure severity. To achieve equally severe seizures, our paradigm involved epochs of 30-s stimulation that were presented for 1 hour and repeated daily.

Following seizure induction, humans and animal models exhibit a post-ictal state characterized by lethargy. The mechanisms contributing to the transition from ictal to postictal state remain elusive: however, candidate neural mechanisms include increased GABA-ergic inhibition, synaptic effects of neuromodulators such as endocannabinoids or adenosine, neuronal exhaustion, neurotransmitter depletion, increased extracellular potassium, and decreased extracellular potassium^69, 70^. In our model, we could not induce a sequential seizure during this state, which typically lasts about 3-6 minutes from the end of a previously elicited seizure (Fig 3e). These data indicate the possibility of a potential mechanism involving resource depletion as opposed to one that might involve a change in cell inhibition. The support for this distinction is that subsequent light stimulation appears to have no additional effects on seizure-related behavior during this refractory period.

We next compared the phenotype in our optogenetics model with the well-established, chemo-convulsant kainic acid (KA) model^71^ (n=4 mice). KA is nondegradable glutamate analog that, when injected intracranially or systemically *in vivo*, induces behavioral seizures and neuronal excitotoxicity^72, 73^. A comparison between optogenetic VPM activation and KA injection revealed a number of common behaviors during severe seizure episodes (supp video 5, Fig 4a). Specifically, both models exhibited tensing of the body (stage 1), hunching posture, erect straub tail, facial and forelimb clonus (stages 2-4), and eventual loss of upright postures (stage 5) with convulsions leading to erratic jumping (stage 6, Table 2). The similar seizure-related behaviors observed in both our *Ntsr1^Cre^;ChR2* model and the KA model argues that a core set of thalamic circuits may be central to seizure initiation. The question we next address is how these behaviors relate to activity.

**Figure 4:**
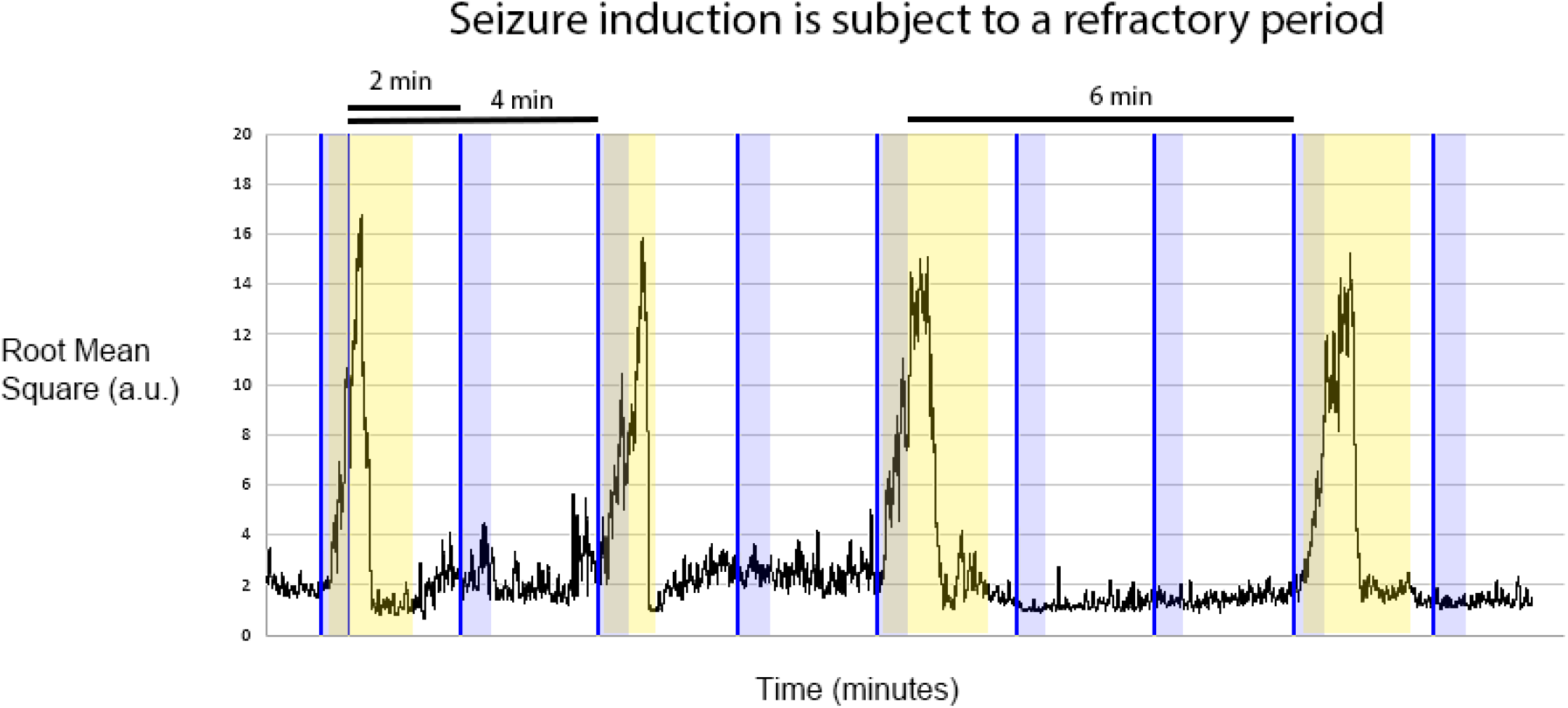
Optogenetic stimulation of the VPM mimics tonic-clonic seizures. (a) Still time-lapse images of the seizure phenotype in the optogenetic (above) and kainic acid (below) models of seizure induction demonstrate parallels in behavior beginning with repetitive facial and forelimb clonus and ending in a loss of posture during convulsions. Arrows point to examples of clonus and an erect, straub tail, both defining features of rodent seizures. (b) Schematic of electrode placement for EcoG (EEG) recordings based on stereotaxic coordinates from the skull landmarks bregma and lambda. Brain overlay adapted from Allen Brain Atlas. (c) EEG signatures during behavioral seizures (boxed region) are synchronous between cortex (blue) and cerebellum (red). This pattern is consistent between optogenetic (above) and kainic acid (below)-evoked seizures.

### Electrocorticography (EcoG/EEG) activity reflects a widespread presence of seizure activity

Electroencephalography (EEG) is an electrophysiological measure used for diagnosing seizure conditions in humans. It is also an important tool for verifying animal models that mimic these conditions. EEG measures population-wide neuronal activity using a series of electrodes typically placed on the surface of the scalp^74^. In mice, increased signal quality is achieved by placing the electrodes directly above the brain. Due to this difference in electrode position, this technique is referred to as electrocorticography (EcoG); for simplicity, we will hereafter refer to both as EEG.

The presence of seizures as a behavioral phenotype in rodents can have components that overlap with other disease-like behaviors. For example, paroxysmal dystonia, a condition characterized by sudden attacks of antagonistic muscle co-contractions, can appear phenotypically similar to rhythmic tonic-clonic seizures^75, 76^. As such, dystonia-like movements and seizure behaviors can be difficult to distinguish in rodents^77^. To confirm the presence of seizures in the *Ntsr1^Cre^;ChR2* mice, we recorded EEG signals from the ipsilateral somatosensory cortex, bilateral motor cortex, and the cerebellum (Fig 4b). We tested whether VPM activation in the *Ntsr1^Cre^;ChR2* mice results in neuronal hypersynchronization. In accordance with the presentation of seizure behaviors, EEG activity in *Ntsr1^Cre^;ChR2* mice showed high amplitude, hypersynchronized activity across the cerebral cortices and cerebellum (Fig 4c, 5a, n=11 mice). We observed similar EEG responses in the KA model (Fig 4c, n=4 mice). This hallmark seizure activity was consistent across all optogenetically-induced mice that exhibited behavioral seizures (supp Fig 5, video 6). Figure 5b shows an example of EEG traces from video-verified seizures in a single mouse during superimposed periods of stage 2 (head nodding and facial clonus) and stage 6 (severe TC seizure with loss of posture and erratic jumping) seizures. In each scenario, the EEG signal reflects the seizure severity at that point in time: during stage 2 clonus, the spike amplitude rose above baseline but was weaker than that during stage 6 convulsions. The presence of high amplitude and hypersynchronized EEG activity and the resemblance of the induced behaviors to an established model provided compelling evidence that our optogenetic paradigm indeed induces TC seizures.

**Figure 5:**
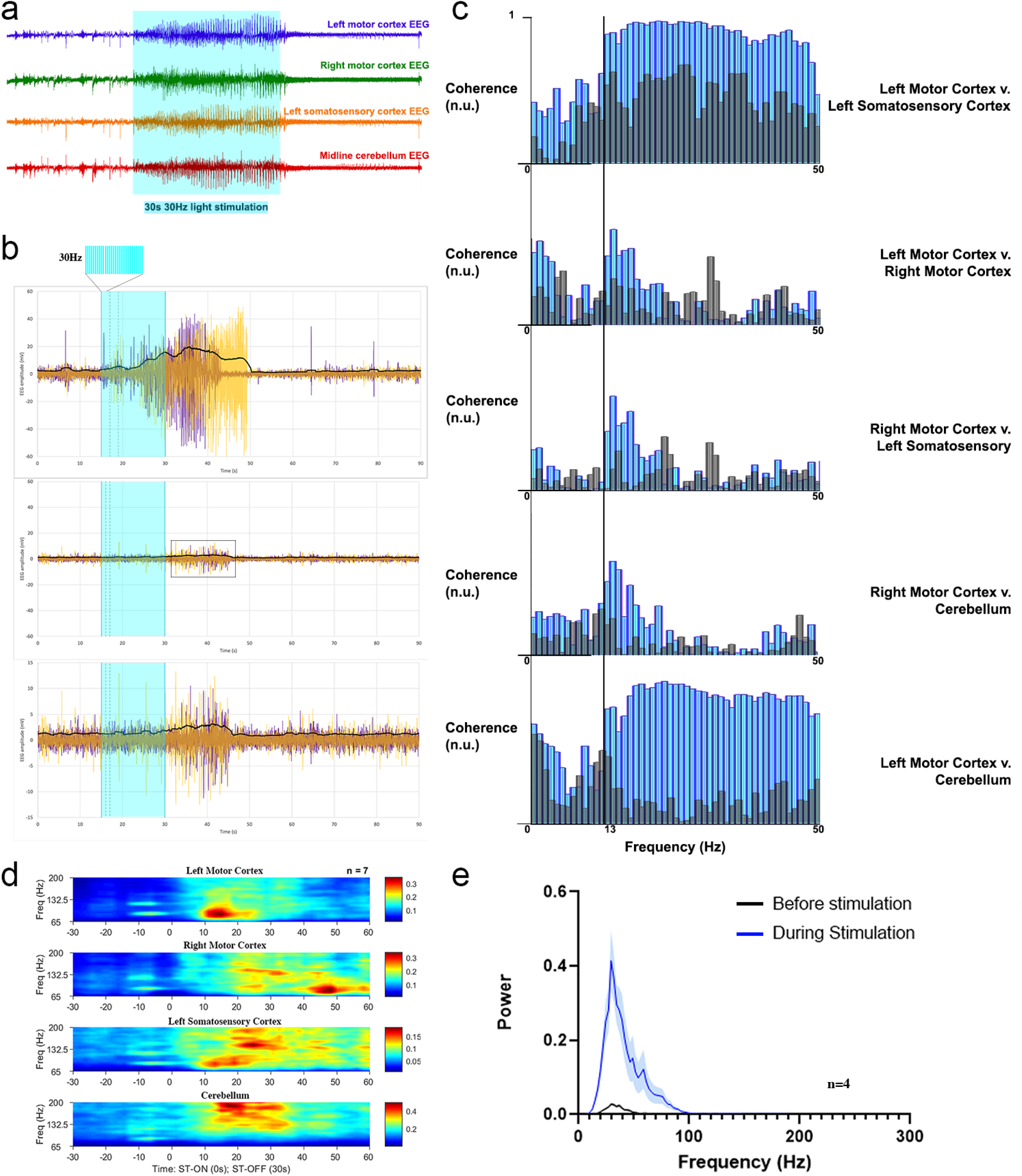
EEG analysis confirms seizure induction. (a) Example of EEG signal from all four recorded brain regions shows high amplitude, synchronous activity during the optogenetic stimulation period (blue box) (b) An example of two superimposed signals (yellow and purple) of averaged EEG (somatosensory cortex, left and right motor cortices, and cerebellum) within a single mouse before, during, and after optogenetic stimulation. Blue shaded region indicates duration of a 30Hz train of light delivered through a fiber optic targeted to the unilateral VPM. Vertical dotted lines demonstrate time at which behavioral seizure is first visualized when both instances are aligned by optogenetic start time. The first plot shows two examples of a stage 6 seizure, whereas the second plot shows two overlaid examples of a stage 2 seizure where only forelimb clonus was observed. The third plot scales the y-axis of the stage 2 seizures to show the change in amplitude from baseline. Black graphed line overlaid on all three plots represents the average root mean square of the EEG signal for both superimposed seizure cases, showing a clear increase from baseline specifically during the behavioral seizure. (c) Example of coherence plots from a single mouse based on the raw EEG signals from four brain regions. The data shows the greatest increase in coherence from before the seizure (gray) to during the seizure (blue) between the left somatosensory cortex/left motor cortex and the left motor cortex/cerebellum. (d) Time frequency heat-plot averaged across mice shows the power of dominant high-power frequencies of EEG shifts from the site of seizure origin (left motor cortex, which is positioned above the VPM) to the contralateral hemisphere (right motor cortex). Time = 0s is aligned to the start of optogenetic stimulation. At 30s, the stimulation is terminated but behavioral and EEG seizure continues independent of the light source and is sustained in cerebellum and left somatosensory cortex. N=7 mice. (e-f) Averaged plot of the power distribution before stimulation (black) versus during stimulation (blue) demonstrate that the most robust changes occur below 100Hz. The same data is shown on a log scale in (f). N=4 mice

### Optogenetic activation of the VPM leads to the coherence of cerebral cortical activity

The synchrony exhibited during seizure-derived EEG occurs as a result of summed dipole fields of contributing neurons in the local electrode region^78^. While it has been argued that this synchrony is driven by only a sub-population of synchronously co-active neurons,^79^ it is nonetheless a consistent measure of seizure activity. Seizure activity is typically highly rhythmic covering a broad spectrum of frequencies^80, 81^. To evaluate the synchronization of rhythmic seizure activity between regions in our model, we examined the coherence of EEG oscillations between the motor cortices, ipsilateral somatosensory cortex, and the cerebellum (Fig 5c, n=4 mice). The left motor cortex and left somatosensory cortices, both having direct neuronal connectivity with, and physically closest to, the stimulation site (left VPM), were most reflective of the intense activity at the seizure origin. Interestingly, these regions exhibited the highest coherence during seizures, particularly in frequencies up to 50Hz (Fig 5c, supp fig 6a). Conversely, the right motor cortex showed a more modest increase in coherence with the left hemisphere, likely due to its position contralateral to the seizure origin. Still, even a lower degree of synchronization supports the generalized spread of the seizure away from its origin, which ultimately incorporates contralateral brain regions into the pathophysiology that eventually drives the behavior.

### The cerebellum and VPM express highly coherent EEG activity during seizures

The cerebellum is typically used as a location of reference for EEG signal collection, an idea based on data showing that the cerebellum exhibits minimal EEG changes during seizure episodes^82–85^. In contrast to these data, we found elevated cerebellar EEG activity^86–89^ that was tightly coupled to the left motor cortex, specifically during seizures occurring between 13 and 50hz (Fig 5c, n=4 mice). The increase in interregional coherence involving the cerebellum was in accordance with our finding that the cerebellum expresses both high amplitude and hypersynchronous activity during optogenetic-induced seizures that were triggered by VPM activation (Fig 4c, 5a, 5c, supp Fig 6c).

### A patterned shift in brain activity emerges as the seizures generalize across regions

EEG can be used to understand, predict, and treat seizures by performing spectral analyses of the signal^90^ and testing for an increase in the power of frequencies between 0.5 and 30Hz. Here, we test whether optogenetic-induced seizures produce a shift in activity by performing wavelet-based time-frequency analyses of the EEG activity. To do so, we examined changes at the following frequency bands: delta (0-4Hz), theta (4-8Hz), alpha (8-13Hz), beta (13-30), and gamma (30-60Hz)^91^ (supp Fig 7, n=7 mice). With a fixed power scale across brain regions, we found that the activation pattern across the brain was inconsistent between mice. We therefore reran the analysis using an optimized scaling for each individual in which a specific brain region’s own maximal power frequencies were graphed. Scale optimization revealed a patterned shift in the dominant activity across all sub-domains from the left hemisphere (left somatosensory and motor cortices, and the central cerebellum) to the right hemisphere (right motor cortex) (Fig 5d, supp Fig 7).

In human epilepsy, high frequency EEG signals that are above 60Hz are thought to include noise due to interference from the skin, skull, and muscle^92^. Bypassing the skin and skull with our electrodes combined with a high sampling rate allowed us to examine these higher frequencies in mice. We were interested in activity in these domains as high frequency oscillations (HFO’s) recorded during epilepsy surgeries have been suggested to have biological relevance in localizing the seizure onset zone^93–95^. Thus, we examined whether changes in activity occurred between 60 and 250Hz in our model. Frequency plots with data averaged over multiple mice revealed a patterned shift in high frequency (HF) power reminiscent of that in the low frequency domains: from the site of origin (left VPM) to the contralateral side of the brain (supp fig 7, Fig 5d, n=7 mice).

We next performed power analyses between 0 and 250Hz to gain a better appreciation for whether neural activity changes on a broader scale (Fig 5e,f, supp Fig 8). Power plot analysis demonstrated that there is a significant bias towards an increase in power in the lower frequency band between 0 and 50Hz during the stimulation period (paired t-test of area under the curve, n=4 mice, average of 4 brain regions each, p<0.0001). In fact, the changes that are represented in the heat plots at higher frequencies are masked in the power plots due to the dramatic differences that occur between 0-50Hz. These data suggest that although HFO’s may help to localize the seizure onset, the largest changes in neuronal oscillation power during seizures indeed occur in the lower frequency ranges.

### Optogenetic stimulation of afferent input to the VPM generates a dynamic, heterogeneous firing pattern in the connected thalamic neurons

We next sought to determine the electrophysiological properties that drive optogenetic-induced seizures in *Ntsr1^Cre^;ChR2* mice. Specifically, we asked how light stimulation of fibers projecting to the VPM affect resident thalamic neuronal activity to initiate seizure behavior. To address this question, we used pulled glass electrodes to record single unit activity in the thalamus of awake, behaving mice implanted with fibers targeted to the ipsilateral VPM. This technique allowed us to examine changes in neuronal activity while simultaneously delivering subthreshold levels of light that are capable of initiating the onset of behavioral seizures but without developing convulsions. To achieve the necessary stability, mice were secured over a freely rotating wheel with a headplate and provided with periods of habituation (Fig 6a). With this approach, we successfully recorded the activity of more than 70 thalamic neurons using an *in vivo* extracellular technique (n=5 mice).

**Figure 6:**
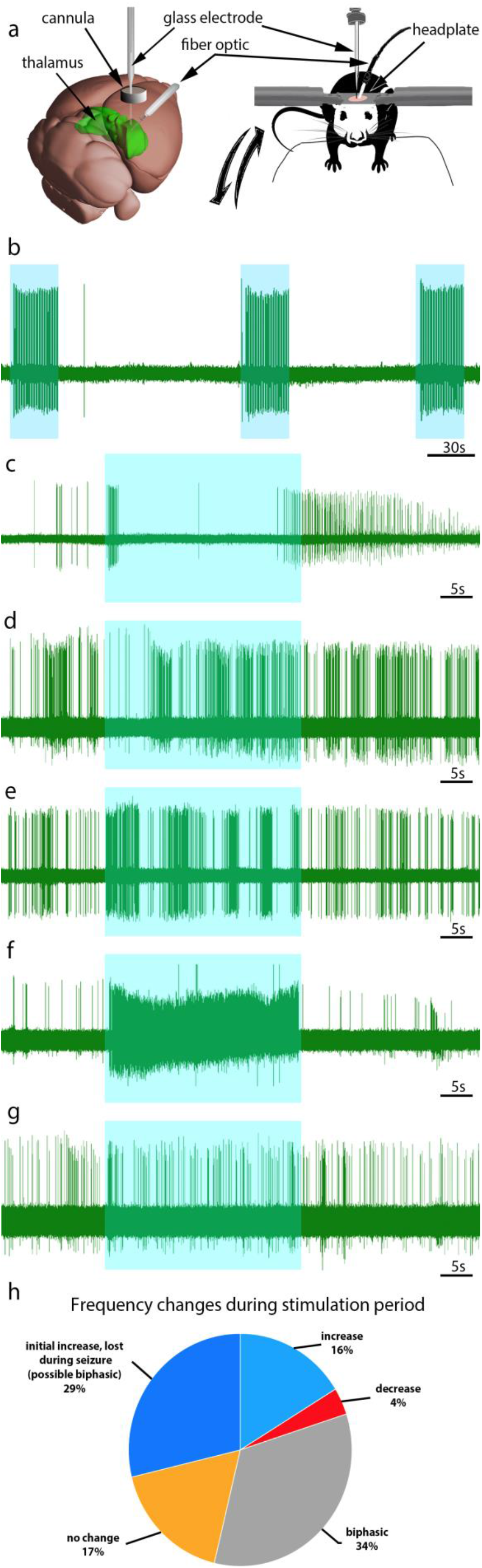
Single cell frequency is altered in the VPM during seizure activity. (a) Schematic of the optogenetic setup. We target the VPM (thalamus, green) with a fiber implanted at an angle and create a craniotomy above the cortex to access the VPM with a glass-pulled electrode for recordings. The mouse itself remains awake and is head-fixed on a rotating wheel, which allows for movement during recordings. (b) Short, subthreshold-seizure stimulations (2 second optogenetic deliveries, blue overlays) elicit reliable entraining of VPM neurons characterized by strict excitation. (c-g) Delivery of seizure-inducing trains of 30Hz light (30 second optogenetic deliveries, blue overlays) elicits heterogeneous responses in thalamic neurons including discreet biphasic (c), repetitive biphasic (d,e), strict excitation (f), and no change in activity during seizures. (h) Quantification of neural responses to seizure behavior, n=5 mice, 71 cells

With the delivery of subthreshold, 2 second light pulses to the ipsilateral VPM, thalamic neurons expectedly demonstrated a repeatable excitatory response (Fig 6b). However, we found that a 30 second seizure-inducing light simulation did not produce a predictable, single response in thalamic neurons. Instead, in addition to excitation (Fig 6b, 6f), they exhibited various biphasic response profiles (Fig 6c-e) or were unaffected by light stimulation (Fig 6g). Most resident neurons exhibited some degree of firing response – only 17% were unaffected by light stimulation (Fig 6h). We postulate that these nonresponsive neurons represent a class of interneurons that do not engage in hypersynchrony^96^ or are not innervated by *Cre-*expressing axons. Interestingly, only 16% of the recorded neurons exhibited purely an increase in frequency. Most cells engaged in a biphasic response, alternating between periods of rapid firing and silence (Fig 6c-e). Due to a high concentration of T-type calcium currents, neurons in the VPM are known to rebound with bursts of excitation following release from hyperpolarization^97–99^. Therefore, during a seizure, thalamic neurons may transition through discrete properties between states of inhibition and excitation.

### Acute optogenetic-induced seizures do not result in cell death, but they do increase dFosB expression on the side of stimulation

Whether acute or chronic seizures cause neuronal death is still largely debated. In several animal models, including electroconvulsive and chemically-induced seizure models (KA, pilocarpine), postmortem analysis of brain tissue provides evidence of programmed cell death^100–105^. However, kindling models have reported tissue expansion following brief seizure induction rather than direct cell loss^106–110^. To examine whether the *Ntsr1^Cre^;ChR2* model involves cell death following seizures, we compared the cell density of H&E stained cells in control mice versus experimental mice that experienced at least one stage-6 seizure. We did not find a significant difference in the number of H&E stained cells in the control animal thalamus versus the seizure animals, indicating that with acute stimulation, optogenetic-induced seizures do not lead to a significant amount of neuronal loss (Fig 7a, unpaired t-test, p=0.4627 n=5 mice per condition, 2-5 tissue sections per animal).

**Figure 7:**
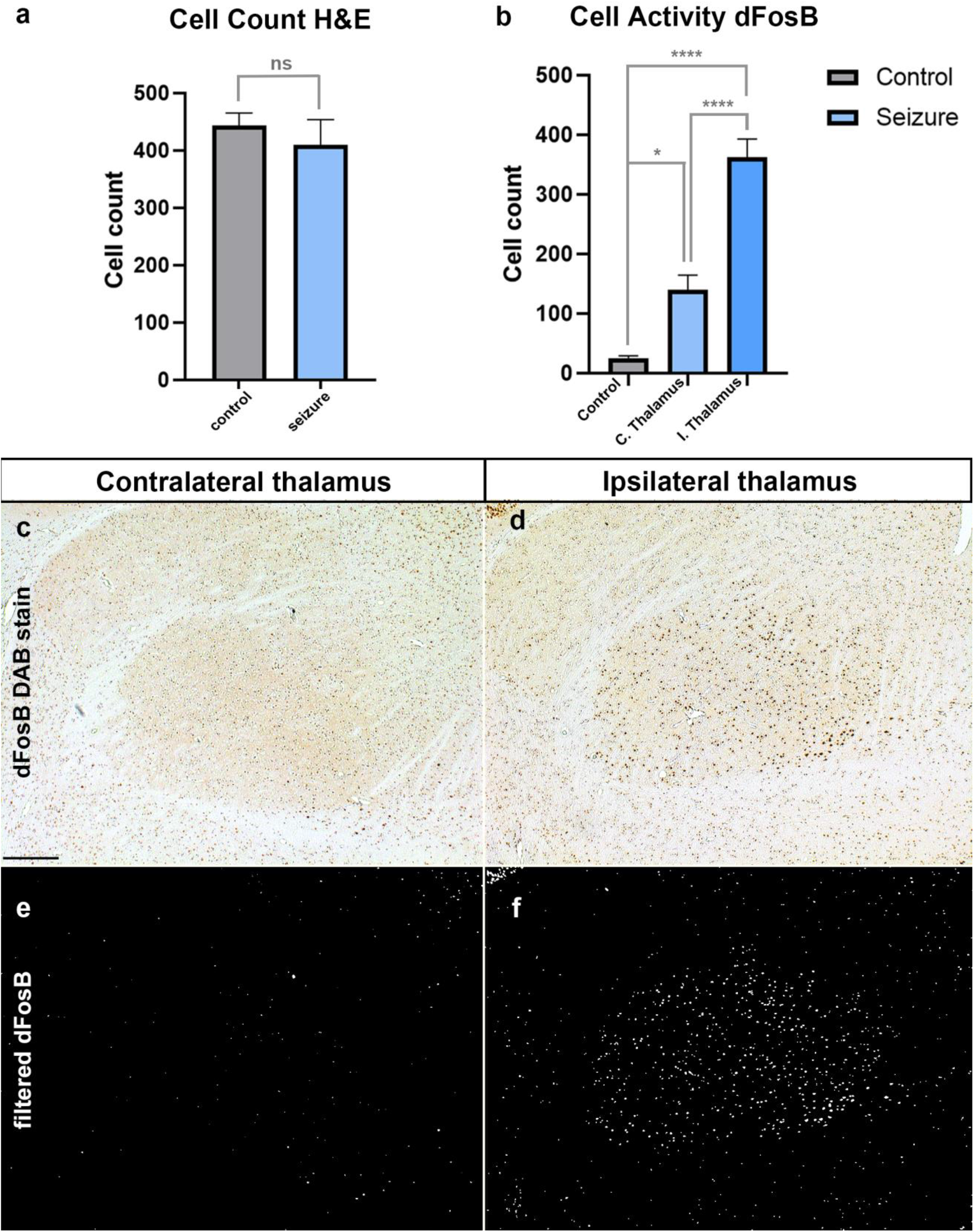
Optogenetic seizure induction does not result in gross cell loss but it does induce a local increase in dFosB expression. (a) Quantification of total cell count using H&E staining demonstrated no cell loss in the thalamus of control versus seizure animals (unpaired t-test, p=0.4627). (b) Quantification of dFosB expression in control animals and seizure animals, both in the ipsilateral thalamus and contralateral thalamus compared to stimulation, demonstrated a graded increase in expression from the contralateral to ipsilateral thalamus. This demonstrates that the most significant changes occurred at the origin of seizure initiation, but also occurred (though to a lesser degree) as seizures spread to the contralateral hemisphere. (Tukey’s multiple comparisons test: control versus seizure contralateral thalamus, p=0.002; control versus seizure ipsilateral thalamus p<0.0001; seizure contralateral versus seizure ipsilateral thalamus, p<0.0001. (c-f) Representative images of DAB visualized immunohistochemical staining of dFosB and their corresponding binary images within the contralateral (c,e) and (d,f) ipsilateral thalamus.

In contrast to cell death, it is well-established that epileptic activity affects long-term neuronal plasticity, which may eventually affect learning and memory^111–115^. Changes in activity are effectively measured by examining the levels of dFosB, a stable immediate-early gene with a long half-life that allows for easy detection^116^. Previous work demonstrates that dFosB is upregulated following seizures, although most studies focus on changes in the hippocampus and cerebral cortex due to their involvement in kindling and temporal lobe epilepsy^117–121^. We therefore tested whether this change in activity is also present in the thalamus, the suspected source of the seizures in our model. We found a significant increase of dFosB in the thalamus ipsilateral to the site of stimulation (Fig 7b-f, Tukey’s multiple comparisons test, p<0.0001, n=4 mice, 3-6 tissue sections per animal). The contralateral thalamus exhibited significantly less dFosB expression than that of the ipsilateral thalamus (p<0.0001), although the level was still higher than in stimulated thalamic nuclei of *Cre* negative littermate controls, (p=0.002). These data indicate that optogenetically induced seizures are accompanied by an upregulation of dFosB in the thalamus, but without significant cell loss.

### Injection of a *Cre*-dependent virus into the VPM delineates source areas of activation

Based on the hypothesis that the VPM represents a strong seizure locus, we next asked which of the *Cre*-expressing brain regions that project to the VPM are capable of eliciting this robust phenotype, or alternatively whether all regions identified above participate in VPM-mediated seizures. We therefore injected retro-rAAV-Ef1a-mCherry-WPRE into the VPM of *Ntsr1^Cre^* mice (Fig 8a, n=5). In this experiment, only *Cre-*positive neurons projecting directly to VPM recombine to express mCherry. After allowing up to 5 weeks for viral tracing and recombination, mice were perfused and prepared for immunohistochemistry. Despite the presence of multiple *Cre*-expressing regions observed in the *TdTomato* cross (supp Fig 1), only the cerebral cortex and cerebellar nuclei demonstrated direct projections to the VPM after virus labeling. The most robust axonal bundles filled by the virus injection originated from the somatosensory areas and cerebellar nuclei (Fig 8b-e). We did not find labeling from other areas that are known to project to the VPM, including the internal segment of the globus pallidus, hippocampus, zona incerta, hypothalamus, and amygdala.

**Figure 8:**
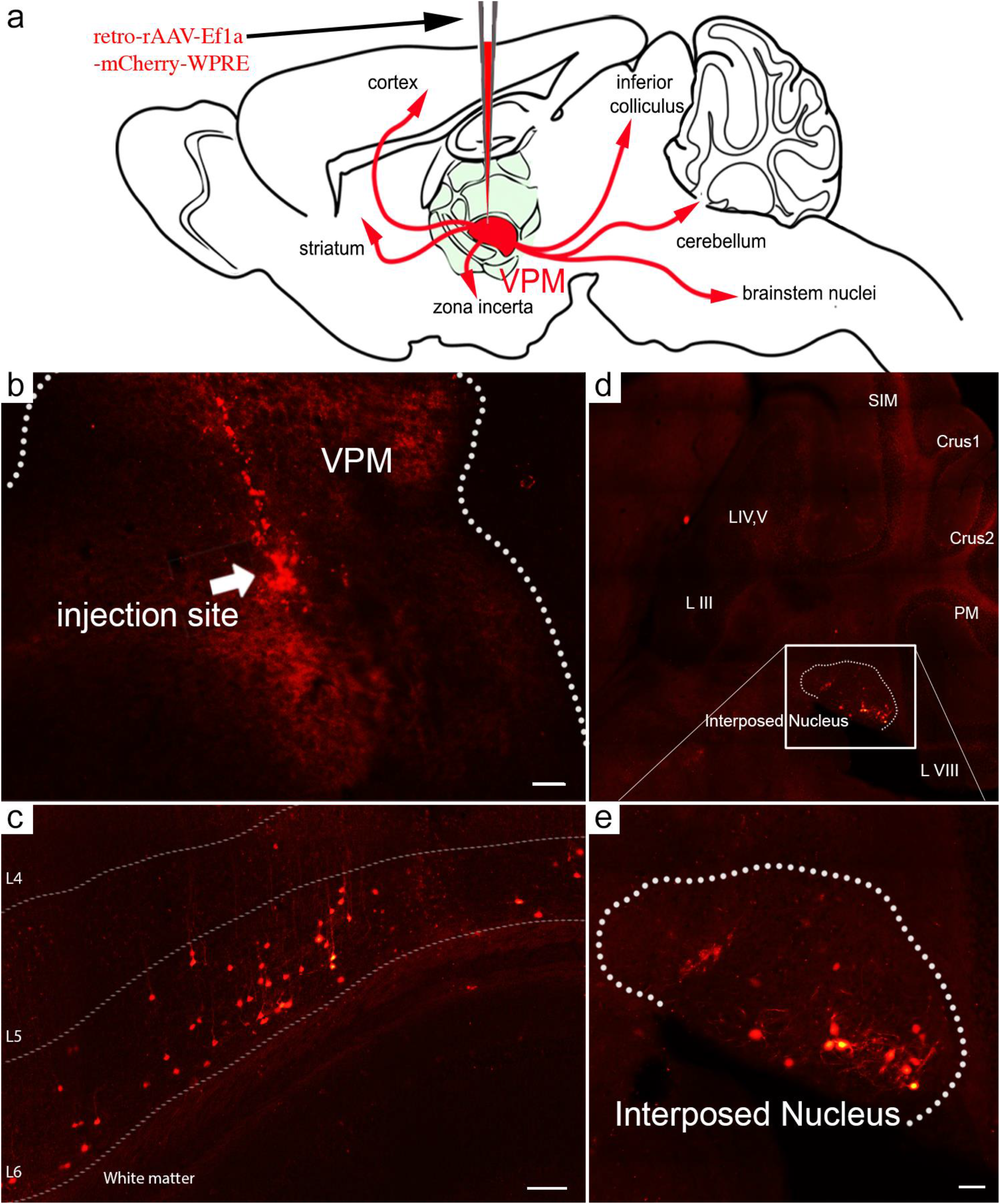
Cre-dependent viral retrograde tracing from the VPM reveals input primarily from the cerebral cortex and cerebellar nuclei. (a) Schematic of the experimental setup. *Ntsr1^Cre^* mice received unilateral injections with varying volumes of pAAV-Ef1a-flex-RUBY ranging from 138nL to 680nL into the left VPM of the thalamus. Virus taken up by local VPM neurons was monosynaptically traced in the retrograde direction to the source neurons that send the projections. In these mice, only the cerebral cortex (b, scale bar = 100um) and cerebellar nuclei (c, scale bar = 50um, d, e, scale bar = 50um) demonstrated a strong marker signal after the injections.

### Drug-induced inhibition of the cerebellum prevents the induction of seizures from the VPM

Based on the virus labeling of the cerebral cortex and cerebellum and that these regions mediate a strong influence on VPM-induced seizure activity, we asked whether both the cerebral cortex and cerebellar nuclei are necessary for initiating the phenotype. To parse apart the relative contribution of each region, we used lidocaine to reversibly block each brain region. Due to the rapid kinetics of lidocaine and the reversibility of its effects, we performed within-trial comparisons of seizure induction before administration, immediately after lidocaine delivery, and after lidocaine washout. We delivered 4% lidocaine through implanted cannulas that bilaterally targeted the cerebellar nuclei (Fig 9a), somatosensory cortex, and motor cortices (Fig 9b). Based on previous studies, we calculated that the neural inactivation window would extend approximately 5-30 minutes after initial delivery^122–125^. Furthermore, previous work from our lab used fluorescent conjugated antibodies to detect the spread of lidocaine and determined that the drug can travel approximately 1mm from the injection site^126^. By delivering lidocaine to each of these regions following seizure induction, we evaluated whether ablation of functional connectivity interrupted the ability to elicit subsequent seizures. We found that pharmacological inhibition of the cerebellar nuclei (Dunnett’s multiple comparisons test, p<0.0001), but not somatosensory (p=0.9562) or motor cortices (p=0.8835), eliminates seizure induction (Fig 9, supp video 5, n=4 cerebellar ablation, n=2 per cortex ablation). Following lidocaine washout, the ability to induce a seizure was rescued. Together, these data support the hypothesis that cerebellar connectivity to the VPM is essential for seizure initiation in this optogenetic model, whereas specific regions of the cerebral cortex may be more heavily involved in generalizing and sustaining the seizure only after it has been initiated elsewhere in the brain.

**Figure 9:**
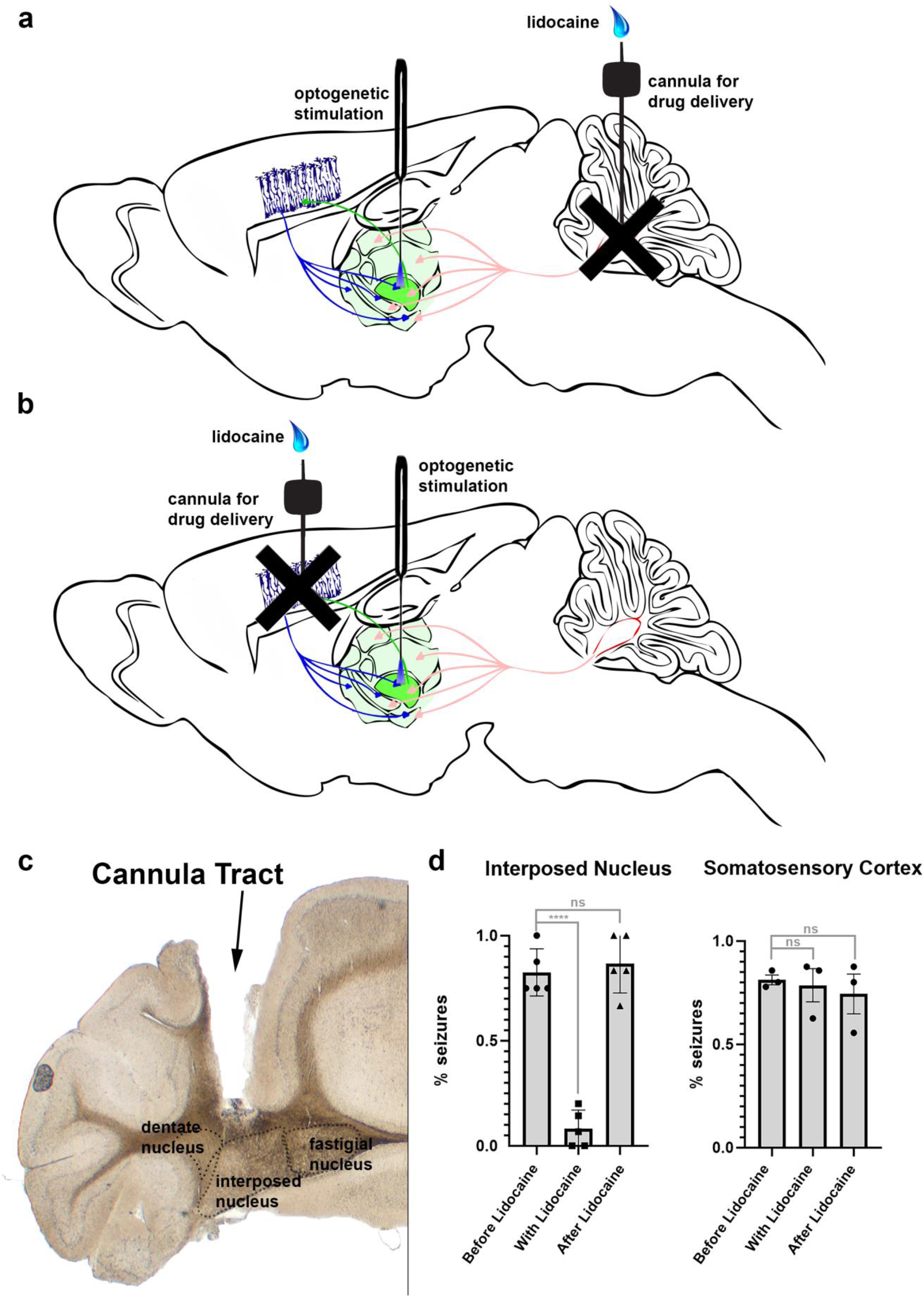
Pharmacological inactivation of individual Cre-expression inputs to the thalamus reveals the requirement of the cerebellum for seizure initiation. (a) Schematic of the experimental setup for cerebellar ablation includes a surgically implanted fiber optic targeted to the VPM and bilateral cannulas targeted to the interposed cerebella nuclei for the purpose of lidocaine delivery. (b) Schematic of the experimental setup for cerebral cortical pharmacological ablation includes a surgically implanted fiber optic targeted to the VPM and bilateral cannulas targeted to the somatosensory or motor cortices for lidocaine delivery. (c) Example of a cannula targeting to the interposed cerebellar nucleus in a 40-um coronal slice. (d) Quantifications of scored seizures after providing trains of 30Hz blue light pulses to the VPM before lidocaine delivery, immediately following delivery, and after lidocaine washout. A significant decrease in ability to elicit seizures occurred following cerebellar interposed nucleus ablation (Dunnett’s multiple comparisons test, p<0.0001), but not somatosensory cortex (p=0.9562) or motor cortical (p=0.8835) ablation.

## Discussion

### Optogenetics provides temporal and cell-type specific regulation to model seizures

There are now many animal models of seizure including mouse, rat, guinea pig, and non-human primate^127–130^. The broad range of models has been advantageous given the heterogeneity of human seizure conditions^131^. However, although each model has its own limitations, their use is nonetheless vital to dissecting circuit mechanisms and other factors that may contribute to the emergence or worsening of these conditions. Currently, there are at least three classes of seizure research models: genetic, chemoconvulsant, and electroconvulsant. While valuable for heredity conditions, those relying upon genetic mutations often do not reflect the cryptogenic forms of epilepsy, which account for over 60% of human cases^132^. Electro- and chemo-convulsant models begin with a healthy brain at baseline, but can introduce secondary effects that may obstruct the analysis of seizure pathophysiology^133–136^. Furthermore, in all three classes of models, the control of seizure initiation can lack temporal precision. We therefore devised an optogenetics approach to overcome these hurdles. We used cell-type specificity to modulate a genetically defined pool of neurons that evoke seizures in otherwise normal mice and then we tracked the spread of abnormal activity. Using an *Ntsr1^Cre^* transgene to drive the expression of a light-sensitive channel in major seizure-engaged regions, we were able to test the anatomical, neuronal, and circuit properties of seizure induction. This model provides control over neuronal activity, an ideal feature for disentangling seizure dynamics. In addition, because optogenetic-induced seizures are under precise temporal control, the stimulation paradigm reliably initiates the seizures within a predictable time window.

The use of optogenetics is not new in the seizure field: a number of studies have benefitted from its advantages to abort seizures in established models of genetic or acquired epilepsy^51, 137, 138–145, 146– 151^, and a few have demonstrated successful induction^138, 152–155^. Current models of optogenetic-driven seizures, some of which exhibit kindling^156^, provide an important and novel platform for studying the specific role of defined neuronal populations in epileptogenesis without cell loss or glial activation. However, our model differs from previous optogenetic models in a few important ways. First, our mouse model does not include viral injections of channelrhodopsin. Although use of viruses can indeed include specificity of cell type targeted via genetic promotors within the viral construct, this extra surgical step can lead to differential targeting of neuronal subsets that lacks consistency from mouse-to-mouse. Second, with accurate fiber optic targeting, our line exhibits severe tonic-clonic seizures upon the first train of light delivery. Current optogenetic models require delivery of light stimulus over a period of days to weeks, drastically increasing experimental time^156^. Third, our model does not include or require the use of seizure-sensitive backgrounds, whether genetic or primed with seizure-inducing drugs. Some optogenetic stimulation paradigms induce convincing TC-seizures within the first few stimulation sessions, but require the use of acute seizure-inducing drugs such as 4-AP with or without the combination of Mg2+ or NMDA^138, 153, 154, 157^. Thus, we greatly simplify and streamline the procedure for optogenetic induced seizure induction using a genetically tagged channelrhodopsin mouse line.

### *Ntsr1^Cre^-ChR2* mice can be used to mark and manipulate ictogenic brain regions

Classical brain regions known to play a major role in seizure pathology are the cerebral cortex and the thalamus. Their roles in seizures are underscored by their reciprocal connectivity with one another and the emergence of neuronal hypersynchrony that can be observed during diverse types of seizures. However, there is now compelling evidence that extra-corticothalamic regions, including the hypothalamus^158, 159^, hippocampus^152, 160, 161^, olfactory cortex^162–165^, amygdala^107, 166–168^, striatum^169–171^, and the cerebellum^54, 172–177^ also play critical roles in seizures. In this study, we used an *Ntsr1^Cre^;ChR2* mouse to test the contributions of the cerebellum and its connections in initiating seizure activity.

The *Ntsr1^Cre^* mouse line has been previously used to demonstrate cerebral cortical involvement in absence epilepsy. Specifically, deletion of *Cacna1a*, which encodes for a P/Q-type voltage gated calcium channel alpha-subunit, in layer VI cortical neurons of *Ntsr1^Cre^* mice generated spontaneous absence epilepsy^178^. Other studies of absence seizures also attribute layer 5/6 cortical neurons in the facial somatosensory cortex as players in epileptic discharge in a rat model of epilepsy^23^. These findings are intriguing as a defining feature of rodent TC seizures, including those elicited in our model, is facial clonus. Our data therefore suggests that absence and TC seizures may have overlapping networks in their pathology, particularly regions such as the somatosensory cortices.

It is interesting to note that the optogenetic-induced seizures that we initiated outlast light delivery, becoming independent of the original stimulation (supp video 4). Combined with the gradual worsening of the overall seizure phenotype with repeated stimulations, this raised the possibility that our model was somewhat equivalent to a kindling paradigm^156, 179–181^; however, we never observed spontaneous seizures in any of our experimental mice (n=24). Nonetheless, we cannot rule out that more subtle seizure-like activity occurred during periods in which the mice were not monitored. We are further convinced that these mice do not experience kindling because given an extended period of multiple days without stimulation, the mice return to baseline seizure severity upon the resumption of stimulation sessions. It is possible, therefore, that while repeated stimulations over the course of hours or days potentiates seizure-involved circuitry, presumably due to potential recruitment of nearby seizure loci, a sufficient length of time between stimulations could allow for recovery of those circuits. This is interesting considering that a single animal can toggle between periods of seizure and behavioral normalcy. We validated our seizure phenotype with EEG. This was important as we did not observe significant changes in the 5-7Hz range (corresponds to 3Hz in humans), a feature used to diagnose absence epilepsy^182^. We conclude that absence seizures are also an unlikely component in our model. Rather, the generalized TC seizure activity exhibited behaviorally can be attributed to a broad increase in low- and high-frequency EEG activity.

### Co-activation of a discrete set of inputs to the VPM accounts for seizure initiation

Neurons within the thalamic nuclei have distinct genetic profiles, morphologies, firing patterns, and synaptic properties ^183–185^. Despite findings that argue for homogeneous activity during seizures, calcium imaging and multiarray studies indicate the presence of heterogenous activity among neuronal populations both at seizure onset as well as throughout the seizure period^96, 186–191^. Furthermore, numerous studies in animal models have established that a proportion of cells within the epileptic brain remain largely normal during seizure episodes. For example, in a cat model of focal epilepsy, only 30% of recorded neurons demonstrated an epileptic response^96^. In humans and monkeys, as many as 51% of neurons recorded by multielectrode arrays were nonresponsive during seizures^187–191^. Here, we demonstrate that perhaps a higher proportion – as many as 80% - of VPM neurons may serve an active role. Whether this is true for other regions has yet to be resolved.

Consistent with the presence of heterogeneous activity, there is now evidence that instead of excitatory neurons being exclusively responsible for initiating seizures, an increase in interneuron activity may first cause local populations of neurons to become quiescent, only to subsequently rebound with sudden bursts of sustained activity^138, 192–194^. It has been suggested that parvalbumin-expressing interneurons may be particularly important for this form of ictogenesis^138, 192–194^. The VPM is known to target both excitatory cortical pyramidal cells as well as GABAergic parvalbumin interneurons^195–197^. In fact, in an investigative long-range tracing study, the authors determined that the VPM accounts for 72% of subcortical input to the PV neurons^198^. In addition, PV-neurons within the reticular thalamic nucleus were reciprocally connect to the VPM^197, 199^. It is possible, therefore, that excitatory-inhibitory neuron interconnectivity between the VPM and surrounding regions partially account for the specific seizure activity observed in our optogenetic induction model.

### Optogenetic-induced seizures induce dFosB expression mainly on the side of stimulation

The immediate-early gene *fosB,* a marker of activity and subsequent plasticity, is altered following seizures^200, 201^. One study found that *fosB-*null mice exhibit spontaneous seizures that resemble the first stage of forelimb clonus seen in our optogenetic model^202^. dFosB, a splice variant of *fosB,* is used to measure changes in activity due to its long half-life. Again, this variant is specifically upregulated in models of epilepsy^117–121^. In addition, dFosB activity has been recently implicated as a component of the learning and memory deficits that occur in Alzheimer’s Disease, and these changes have been attributed to seizure activity in these animal models and human patients^203^.

In our model of VPM-induced seizures, we demonstrate that dFosB expression is increased in the thalamus on the side of stimulation (Fig. 7). Upregulation of dFosB also occurred in the contralateral thalamus, though to a lesser degree. Our interpretation is that the largest change in activity occurs at the seizure epicenter and then spreads to the contralateral hemisphere. Overall, this marker of activity is not affected by light delivery: rather, dFosB expression only increases in the thalamus when a seizure-evoking optogenetic train of light successfully induces seizure activity, as demonstrated by ChR2^-/-^ control animals that received the same light stimulation without experiencing seizures. Previous studies of corneal stimulation to induce TC seizures in rodents observed similar dFosB increases, although in the amygdala and hippocampus^204^. We did not see significant changes in CA1, CA3, or dentate gyrus, nor in amygdala, perhaps due to the different method of seizure induction. Lastly, while previous studies have reported differences in cerebellar activity through decreases in dFosB expression in the cerebellar nuclei^172^, we did not observe such changes. This is likely a result of our experimental paradigm, which involves only one week of seizure inductions, with likely correspondence between the models if a chronic paradigm was used.

### Cerebellar connectivity is required for tonic-clonic seizure initiation at the VPM

Cerebellar contribution to seizures is one with a long-standing history: as the first deep brain stimulation target for treatment of epilepsy in the 1970’s, the cerebellum was initially a major site of focus in seizure studies. Evidence for a clinical role of cerebellum in epilepsy is abundant: patients with tubercular abscesses in cerebellum have been reported to experience seizures. Likewise, those with gangliomas in the cerebellum noted elimination of seizures upon their resection^36, 205^. In patients with histories of repeated seizure, a correlation between the number of seizure episodes and cerebellar atrophy through Purkinje cell death is noted through MRI imaging studies^206, 207^. Further evidence of cerebellar involvement lies in epilepsy comorbidities: in many cases, patients with seizure disorders present with known cerebellar-derived comorbidities including ataxia, dystonia, and tremor^208, 209^. Recently, studies have shown reliable eradication of the physical symptoms and abnormal EEG with treatment targeted to the cerebellar nuclei^49, 51^. EEG and single unit electrophysiology studies corroborate these data by demonstrating robust modulation of neuronal activity in the cerebellum during seizure episodes in various mouse models^210–212^.

Our data demonstrates that the cerebellum may in fact also engage in chronic network hyperactivity during seizures, which supports a growing literature supporting a dynamic role of the cerebellum in epilepsy^19, 43, 44, 47, 50–53, 173, 205, 211, 213, 214^. Based on our time-frequency analyses, we tracked the temporal spread of seizure activity from the left motor cortex to the left somatosensory, cerebellar, and eventually right cortical areas. According to these data, the cerebellum is activated prior to seizure spread and the greatest peak power across frequency bands was in the cerebellum (supp Fig 8). It is interesting to note that cFos expression is abundantly activated in granule cells in epilepsy^173^. In cases of cerebellar gangliomas where seizures consequently manifest, resection of the tumor eliminates the seizures^36^. Similarly, clinical studies note the presence of cerebellar gangliomas in cases where the sole behavioral symptom is severe seizures^215^. Ming, et al. (2021) hypothesize that cerebellar disinhibition leads to cortical activation, thereby exacerbating or initiating seizures^216^. Our data support the hypothesis that cerebellar activation can drive seizures.

However, there is also evidence proposing the opposite: that decreasing cerebellar activity worsens seizures. For example, clinical observations since the early 1800’s have noted cerebellar atrophy in patients with prolonged seizures; immunohistochemistry experiments report lower dFosB expression in cerebellar nuclei following seizures, indicating a decrease in neuronal activity^172, 217^; pharmacological inhibition of the cerebellar nuclei increased abnormal EEG activity in absence mouse models, whereas excitation blocked their occurance^50^. Despite these seemingly contradictory results, both scenarios could co-exist. Depending on the pattern of cerebellar activity, it could either synchronize or disrupt downstream networks to modulate seizures. In genetic models of epilepsy, cerebellar firing is abnormal^172, 213, 218, 219^. During periods of absence seizure episodes, the cerebellum switches from rhythmic to asynchronous firing.^50, 219^ This rhythm of cerebellar output may force synchrony between regions like the thalamus and cerebral cortex. In these disease models, where baseline cerebellar activity is already disrupted, delivery of stimulation through DBS or optogenetic pulses of light may disrupt synchronized networks and terminate seizures. However, in models with normal baseline cerebellar activity, delivery of a patterned train of light to cerebellar nuclei instead forces the same bursty pattern that is inherent in epilepsy models, thereby initiating synchrony in upstream regions to cause a seizure. In patients with genetic forms of epilepsy, the cerebellum may therefore be an important target for treatment to correct abnormal output. In idiopathic or cryptogenic epilepsies, which make up an overwhelming 65% of cases, abnormalities in the cerebellum may be at fault for the spontaneous emergence of seizures. In both cases, the cerebellum should be considered as a key player in epilepsy pathophysiology.

## Conclusion

In these studies, we devised an optogenetics mouse model to demonstrate that 30Hz coactivation of cerebral cortical and cerebellar input to the VPM induces severe TC seizures. With viral tracing and reversible inactivation, we show that of these circuit inputs, the cerebellum has a predominant role in seizure initiation, supporting recent work that links cerebellar connectivity to epilepsy. Furthermore, using single-unit *in vivo* electrophysiology recordings performed in seizing mice, we uncover heterogeneous, biphasic activity of thalamic neurons during seizures. Together, these data suggest that cerebellar input to the VPM is a critical source of abnormal signals at seizure onset.

## Acknowledgements

This work was supported by funds from Baylor College of Medicine (BCM) and Texas Children’s Hospital. RVS received support from The Hamill Foundation and the National Institutes of Neurological Disorders and Stroke (NINDS) R01NS089664, R01NS100874, and R01NS119301. RVS is also supported by a grant from the Dystonia Medical Research Foundation (DMRF). RVS, DHH and YL are supported by National Institute of Mental Health (NIMH) R01MH112143. Research reported in this publication was supported by the Eunice Kennedy Shriver National Institute of Child Health & Human Development of the National Institutes of Health under Award Number P50HD103555 for use of the Cell & Tissue Pathogenesis Core and the Neuroconnectivity Core. The content is solely the responsibility of the authors and does not necessarily represent the official views of the National Institutes of Health. JB received support from F31NS115432.

## Contributions

JB, JO-G and RVS designed the experiments. JB, JO-G, TL, and RVS performed the experiments. JB, BB, YL, DHH, and RVS designed code for electrophysiology analysis. JB, JO-G, BB, YL, DHH, BRA and RVS analyzed the data. JB and RVS wrote and edited the paper.

## Ethics

All mice were housed in an AALAS-accredited facility on a 14hr light cycle. Husbandry, housing, euthanasia, and experimental guidelines were reviewed and approved by the Institutional Animal Care and Use Committee (IACUC) of Baylor College of Medicine (protocol number: AN-5996).

## Methods

### Animals

To develop transgenic mice with restricted TdTomato or Sun1 expression, we crossed heterozygous *Ntsr1^Cre^* males (*B6.FVB(Cg)-Tg(Ntsr1-cre)GN220Gsat/Mmucd,* University of California, Davis MMRRC, Mouse Biology Program) to het- or homozygous *ROSA^lox-stop-lox TdTomato^* or *ROSA^lox-stop-lox-Sun^*^1^ lines *(B6;129-Gt(ROSA)26Sor^tm^*^5^*^(CAG-Sun^*^1^*^/sfGFP)Nat/J, Jackson^* Laboratory, Bar Harbor, Maine). These offspring, whose genotype was confirmed through tail snips or ear punches, were used for immunohistochemistry. To gain light-sensitive control over these neurons, we crossed *Ntsr1^Cre^* males to *ROSA^lox-stop-lox-ChR^*^2^*^-YFP^* females. Again, their genotypes were confirmed through tail and ear clippings, but also through immunohistochemistry after experimentation. We bred the mice using timed pregnancies, and designated noon on the day a vaginal plug was detected as embryonic day (E) 0.5 and the day of birth as P0. For experimental studies, mice from P60 to P180 were used. Mice of both sexes were studied, and opsin or *Cre*-negative littermates were used as non-expressing seizure controls. The mice were housed on a 14h/10h light/dark cycle. All animal studies were carried out under an approved institutional animal care and use committee animal protocol according to the institutional guidelines at the Baylor College of Medicine.

### Surgery

Surgery for awake optogenetics was performed as detailed in Ung and Arenkiel, 2012^8^. Briefly, mice were anesthetized with 4% isoflurane in an induction chamber until unresponsive to the toe-pinch reflex. After transferring to a stereotaxic apparatus, isoflurane was lowered and maintained at 2% for the duration of surgery. Using aseptic technique and a sterile work field, the mouse was cleaned and prepped, the skin above the skull opened, and a small craniotomy of approximately 1mm in diameter made above the VPM (AP −1.6mm, ML +/-1.55mm). A polished optic fiber previously glued into a ceramic ferrule (Thorlabs, Newton, NJ, USA; #CFLC230-10) was then gently lowered to the approximate depth (2.8mm) and secured with Metabond Adhesive Luting Cement. For most surgeries in which EEG was also incorporated, 6 additional craniotomies were made above the super colliculi (reference and ground electrodes, AP - 5.21mm, ML +/-1mm), primary sensory cortex (AP +1mm, ML +3.5mm), primary motor cortices (AP −1mm, ML +/-1mm), and cerebellum (AP −6mm, midline). An EEG head mount compatible with a detachable preamplifier (Pinnacle Technology, Inc, Lawrence, KS, USA; #8406) was equipped with six silver ball-tipped wires (A-M Systems, Sequim, WA, USA; #785500) soldered to individual pins. This head mount was glued to the most rostral portion of the skull. The wire free ends were inserted into the craniotomies between the skull and dura. All wires were secured with UV epoxy followed by Metabond Adhesive Luting Cement. Next, the fiber optic implant as well as an EEG head mount were wrapped in dental cement. After the surgery, the mouse was placed in a warming box (V500, Peco Services Ltd., Cumbria, UK) to prevent hypothermia while the anesthesia wears off. Once the mouse was awake and mobile, it was returned to the home cage. The mouse was allowed to recover for 2-3 days before optogenetic stimulation and recordings were initiated.

### Optogenetics and EEG collection

To induce seizures in this mouse model post-surgery, a fiber optic cable powered by a 465nm light box (ALA Scientific Instruments Inc, Farmingdale, NY, USA) was carefully inserted into the unilateral fiber implant surgically targeted to thalamus (VPM). When EEG was included, a preamplifier (Pinnacle Technology, Inc, Lawrence, KS, USA; #8406) was connected to the implanted EEG head mount. All EEG and optogenetic stimulation parameters with subsequent data was recorded through Spike2 software and delivered using a CED Power1401 data acquisition interface (CED, Cambridge, UK). Recordings were paired with live video acquisition and were monitored for 5 minutes before stimulation, the duration of stimulation, and 5 minutes after stimulation. To entrain the circuits to seizures, 1 hour of 30Hz trains of optogenetic stimulation, 2 minutes light on and two minutes lights off, was applied daily in the early afternoon for one week. Depending upon the precise targeting of the fiber optics, severe seizures would typically be triggered upon the first stimulation of the day beginning on day 2-3 if not on the first day. Seizures were reliably induced every day after the first successful induction.

### Tissue processing and immunohistochemistry

The mice were perfused with 1 M phosphate-buffered saline (PBS) and 4% paraformaldehyde (PFA). The brain was then carefully dissected and submerged in 4% PFA for 1-2 days. Following serial sucrose protection (15-30% sucrose in PBS), samples were submerged in OCT and immediately frozen at −80C. For immunohistochemistry, free-floating sections cut on the cryostat were incubated in antibodies in a solution of 10% normal goat or donkey serum and 0.01% Tween-20. For DAB staining, sections were first bathed in H2O2 to quench endogenous peroxidases and then washed 3 times in PBS for 5 min each. For all stains, sections were blocked for 2 h with the blocking solution at room temperature and then incubated in primary antibodies overnight at room temperature. Sections were then washed with PBS 3 times for 5 min each and then incubated in secondary antibodies for 2 h. Sections were washed again 3 times in PBS for 5 min each and mounted onto slides. DAB slides were air-dried overnight and then dehydrated in ethanol before cover-slipping. Slides with fluorescent signal were immediately cover-slipped using FluoroGel as a medium. Antibodies used for immunohistochemistry (primary antibodies) were: calbindin (Rb and mouse (Ms), Swant, 1:10,000); NeuN (Rb, Millipore, 1:500); VGLUT2 (Rb and Gp, Synaptic Systems; 1:1,000); VGLUT1 (Ms and Gp, Synaptic Systems, 1:1,000); NFH (Ms, Covance, 1:1,500); TH (Rb, Millipore, 1:500); GFP (Ch, Abcam, 1:1,000); parvalbumin (Rb, Swant, 1:1,000); calretinin (Rb, Swant, 1:500); Secondary antibodies were: for the DAB reaction, we used horseradish peroxidase-conjugated goat anti-rabbit and goat anti-mouse secondary antibodies (diluted 1:200 in PBS; DAKO); staining for fluorescent immunohistochemistry was performed using donkey anti-mouse, anti-rabbit or anti-guinea pig secondary antibodies conjugated to Alexa 488, 555 or 647 fluorophores (Invitrogen), all diluted to 1:1,500. Photomicrographs of tissue sections were captured using Zeiss AxioCam MRm (fluorescence) and AxioCam MRc5 (DAB-reacted tissue sections) cameras mounted on a Zeiss Axio Imager.M2 microscope or on a Zeiss Axio Zoom.V16. Images of the tissue sections were acquired and analyzed using either Zeiss AxioVision software (release 4.8) or Zeiss ZEN software (2012 edition). After imaging, the raw data were imported into Adobe Photoshop CS5 and corrected for brightness and contrast levels. Schematics were drawn in Adobe Illustrator CS5 and then imported into Adobe Photoshop CS5 to contract the figure.

### Basic histology: H&E staining

For H&E staining, tissue sections were mounted on a polarized slide and dried overnight. Next, the slide was submerged in xylene for 5 minutes. To then rehydrate the tissue, the sections were placed in serial dilutions of ethanol for 2-5 minutes as follows: 100% ethanol, 95% ethanol, 70% ethanol, DI water. Following this rehydration step, slides were placed in hematoxylin staining for approximately 1 minute followed by a water bath for several rounds of washing. Finally, tissue was placed in an eosin stain for 1 minute, rinsed in a water bath, and dehydrated in 70%, 95%, and three rounds 100% ethanol followed by xylene. Slides were then coverslipped with Cytoseal Mounting Media (Fisher Scientific) and left to dry overnight.

### In vivo electrophysiology

For head-fixed, awake recordings, mice were implanted with custom-made headplates and a craniotomy made above the VPM. A fiber optic was also implanted at a 45-degree angle to target the VPM underneath the headplate. After 72 h of recovery, the mice were trained for 30 min per day in a head-fixed apparatus for 3 days before recording. Neuronal spikes were recorded and categorized based on standard stereotaxic coordinates measured from bregma and firing pattern. We examined a total of 70 units in 5 animals. All recordings were collected using ∼10-20MΩ pulled glass pipettes filled with 0.9% saline. The recorded signals were digitized at a sampling rate of 5000Hz into Spike2 (CED, England) where single units were verified with principal components analysis.

### Quantifications and Analyses

All data was analyzed and graphed in Prism9.1.0 and p values were considered significant at values less than 0.05. For differences in relative fluorescence, thalamic nuclei were segmented in Photoshop and their mean gray values, area, and integrated densities were calculated. A region of layer 6 cortex was used as reference for each slice. Data was then transferred to Prism, graphed, and Dunnett’s multiple comparisons test performed with the VPM region used as a control for comparison. All EEG data was collected in Spike2 and analyzed in MATLAB (code available upon request), except in the case of power analyses which were exported from Spike2 and plotted in Prism. Statistics were computed using paired t-tests on area under the curve. Cell count (on H&E stained sections) and dFosB quantifications were performed by first thresholding, converting to binary images, and automatically counting positively stained cells in ImageJ before transferring data to Prism for unpaired t-tests. For lidocaine experiments, presence of seizures was visually scored as 1 (seizure) or 0 (no seizure) for a train of 30 second, 30-Hz stimulations before lidocaine administration and beginning 5 minutes after administration. For washout experiments, mice were returned to their home cages and re-evaluated with optogenetic stimulation and visual scoring 1 hour after initial lidocaine administration. These data were then plotted and analyzed via Dunnett’s multiple comparisons test in Prism.

### Data availability

Data generated in the experiments presented in the current study are available from the corresponding author upon request and also available in the data tables provided.

---

**S1:**
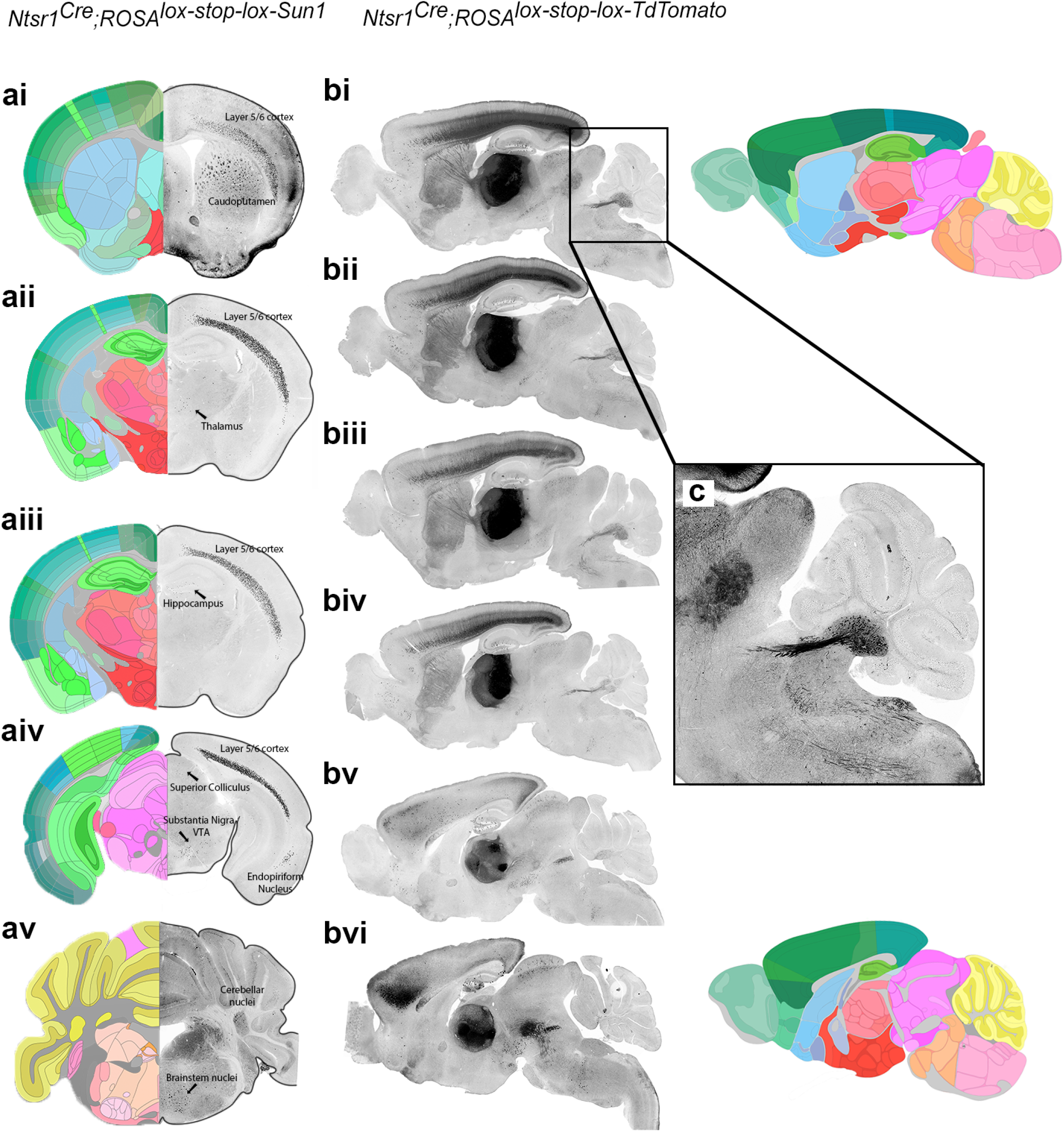
Whole brain expression of the Ntsr1^Cre^ transgene. (ai-av) Coronal sections of *Ntsr1^Cre^;ROSA^lox-stop-lox-Sun^*^1^*^-GFP^* mice show nuclear envelope staining primarily in cortical layer 5/6, the caudoputamen, and the cerebellar nuclei. However, there is also noticeable expression in the substantia nigra/VTA and brainstem nuclei. Although there are some *Ntsr1^Cre^* positive neurons in the hippocampus and medial thalamic nuclei, these are sparse, few, and unlikely to affect optogenetic stimulation paradigms. (bi-bvi) Sagittal sections cut through *Ntsr1^Cre^;ROSA^lox-stop-lox-TdTomato^* mouse brains highlight the robust fluorescence in thalamic nuclei stemming exclusively from incoming axons and fibers of passage, one example of which is seen in panel (c), a dense bundle of *Ntsr1^Cre^* positive axons exiting the cerebellum. Segmented atlas images (adapted from Allen Brain Atlas) are included for reference.

**S2:**
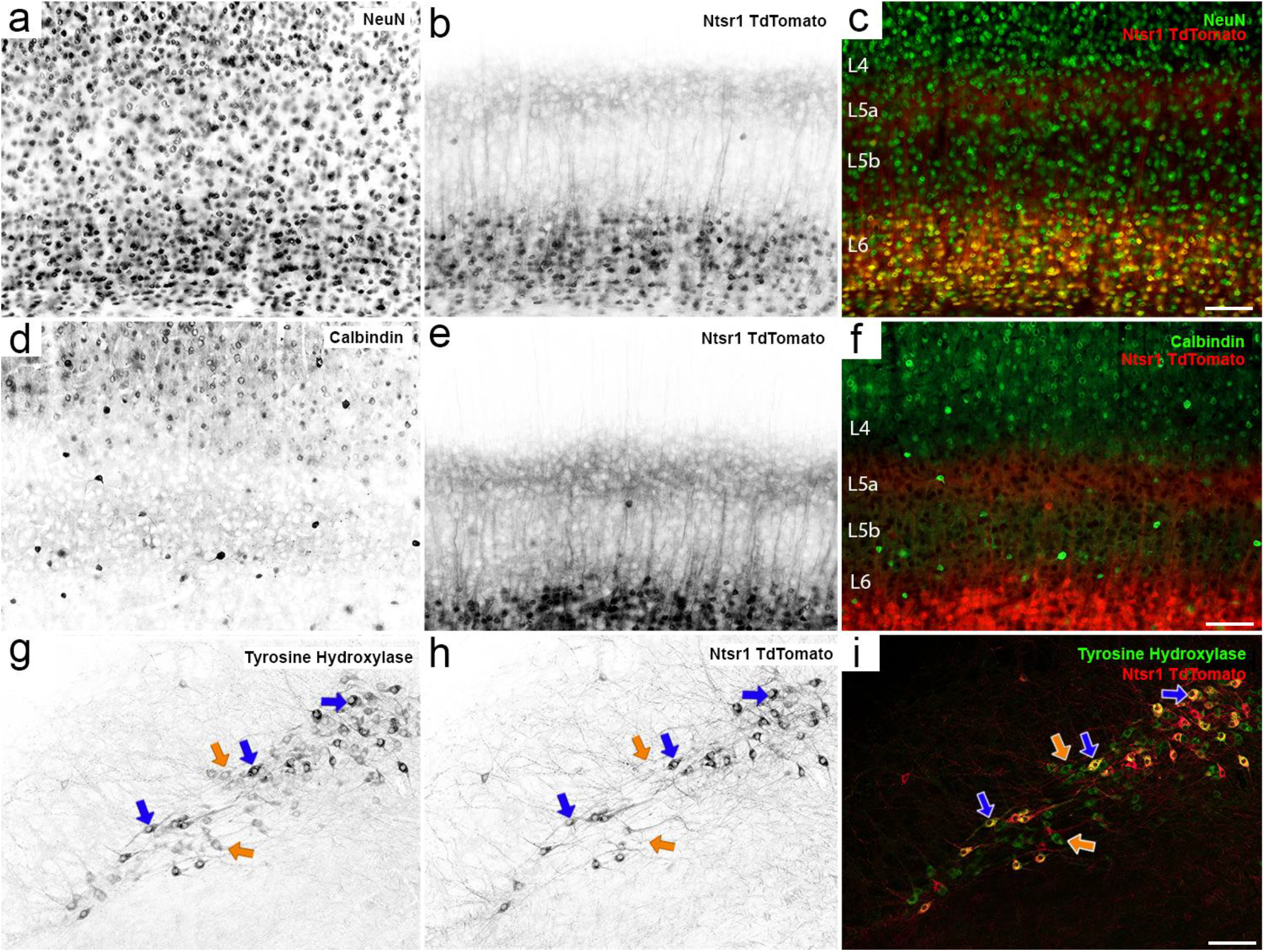
Characterization of Ntsr1^Cre^ positive neurons in the cortex and VTA. (a-c) Co-staining of NeuN in the cerebral cortex of *Ntsr1^Cre^;ROSA^lox-stop-lox-TdTomato^* mice demonstrate specificity of TdTomato signal to layer 6 cell bodies. Scale bar = 100um. (d-f) Calbindin staining, which labels nonpyramidal cells in the cortex, does not colocalize with TdTomato positive cells. Scale bar = 100um. (g-i) In the VTA, TdTomato-positive cells colocalize with tyrosine hydroxylase (TH), indicating that they are dopaminergic (white arrows). However, not all TH-positive cells are TdTomato positive (orange arrows), indicating that the *Ntsr1^Cre^* allele is not expressed in all VTA neurons. Scale bar = 50um.

**S3:**
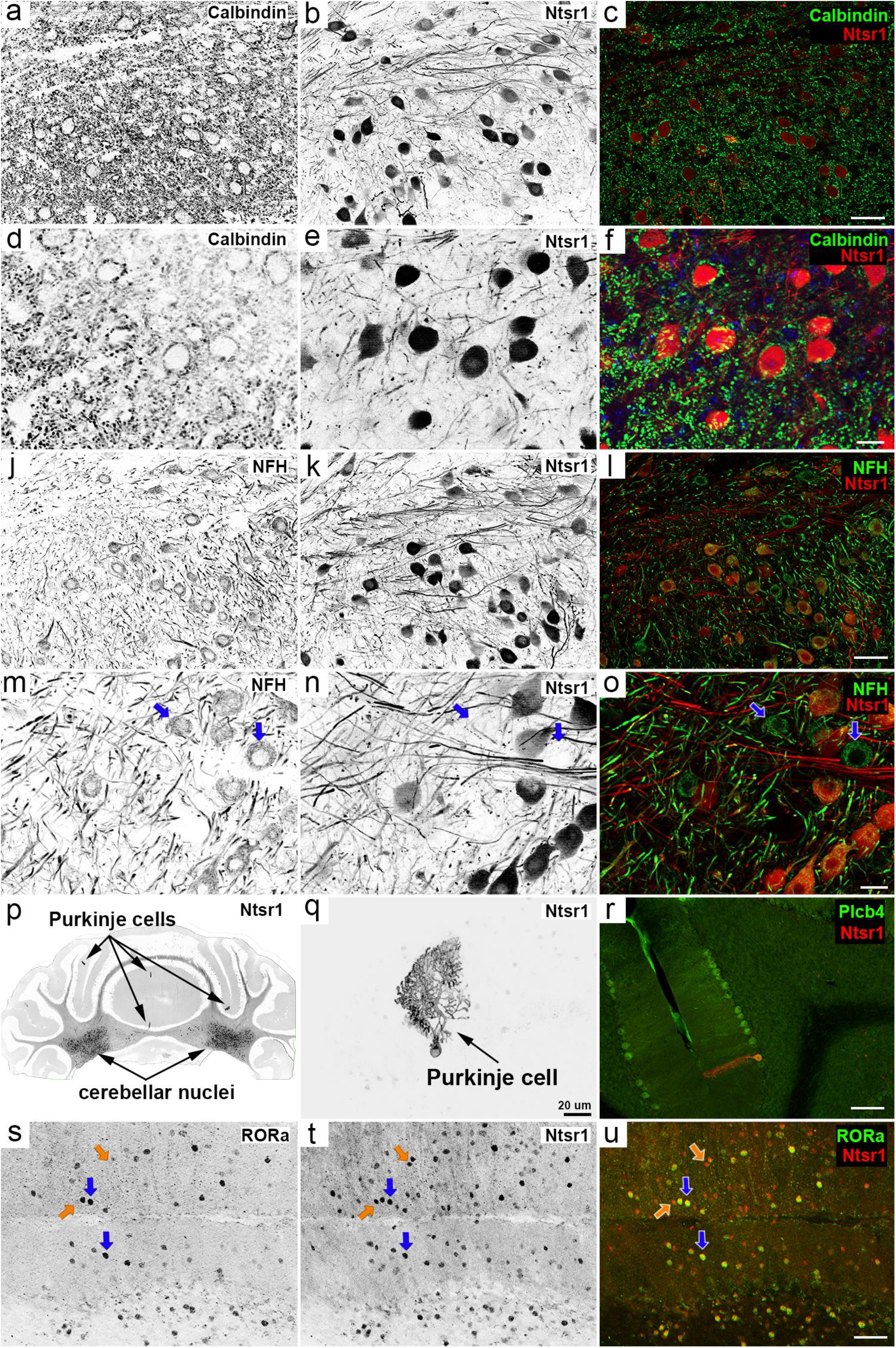
Ntsr1^Cre^ expression in the cerebellum is localized to excitatory cerebellar nuclei neurons, scattered Purkinje cells, and a small population of molecular layer interneurons. (a-f) Calbindin staining (green) demonstrates the relationship between GABAergic projections from Purkinje cells to *Ntsr1^Cre^*-positive cerebellar nuclei neurons (red). Scale bars are 50um (c) and 20um (f). (j-o) Neurofilament heavy chain (NFH, green) colocalizes with TdTomato neurons in the interposed and dentate nucleus, identifying these cells as excitatory projection neurons. Scale bars are 50um (l) and 20 um (o). (p) *Ntsr1^Cre^;TdTomato* coronal section of the cerebellum shows sparse labeling of individual Purkinje cells (arrows). (q) A single *Ntsr1^Cre^*-positive Purkinje cell shows intense expression in the soma and throughout the dendritic tree. Scale bar = 20um. (r) *Ntsr1^Cre^* positive Purkinje cells also express the Purkinje cell marker Plcb4. Scale bar = 100um. (s-u) RORa staining in lobules IX and X demonstrates that some (blue arrows), but not all (orange arrows), molecular layer interneurons (basket cells and stellate cells) express the *Ntsr1^Cre^*-transgene. Scale bar = 100um.

**S4:**
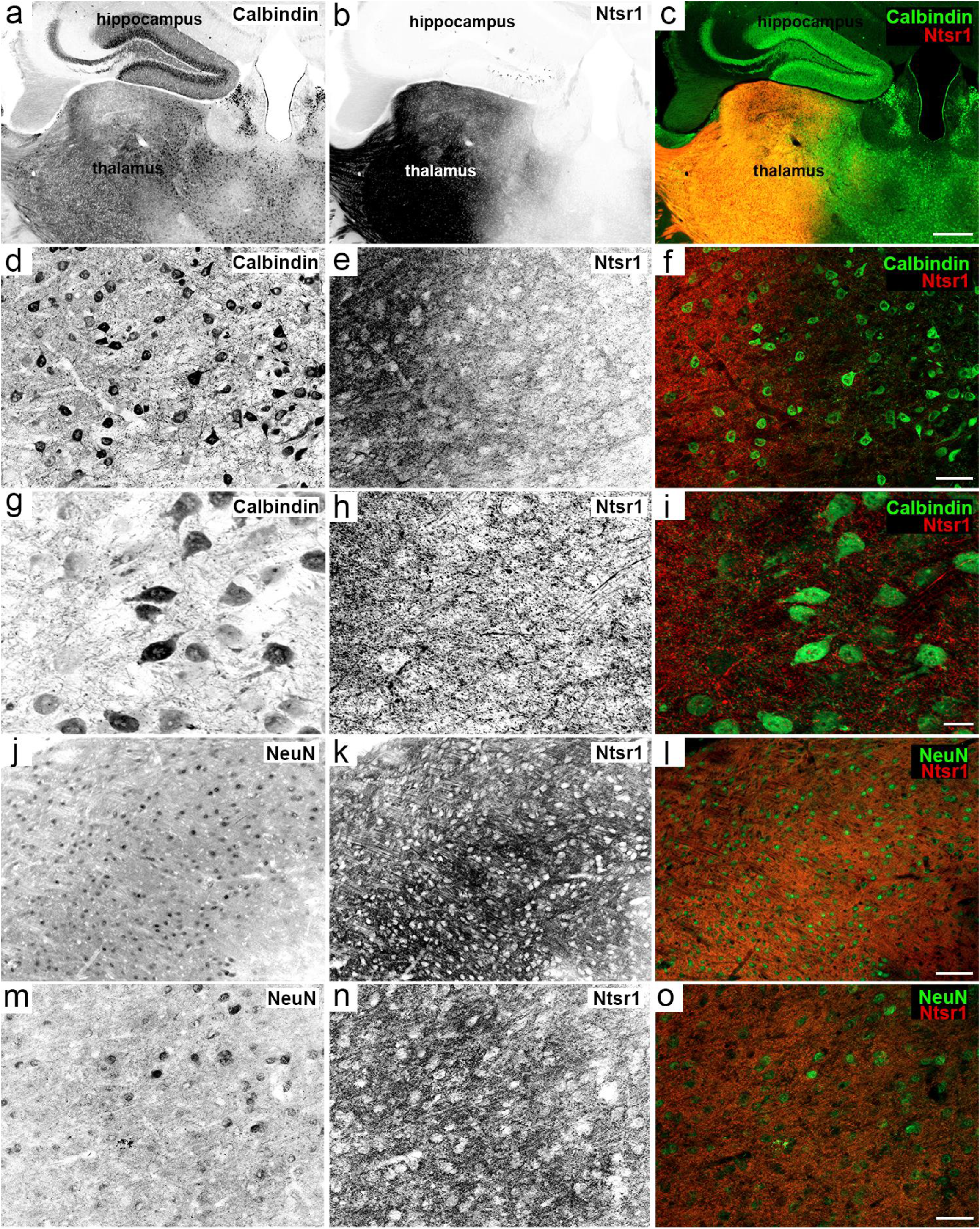
Ntsr1^Cre^ is not expressed in cell bodies of thalamic neurons. (a-i) Serially magnified images of calbindin staining in the thalamus, which is differentially expressed in various thalamic nuclei, demonstrates no colocalization between TdTomato (red) and calbindin-positive (green) thalamic neurons. Scale bars are 500um (c), 50um (f), and 20um (i). (j-o) Staining with a pan neuronal marker, NeuN, does not colocalize with TdTomato signal in the thalamus. Instead, *Ntsr1^Cre^* is expressed in the space around resident thalamic cell bodies (k, n) as local axons or fibers of passage originating from other *Cre*-positive brain regions. Scale bars are 100um (l) and 50um (o).

**S5:**
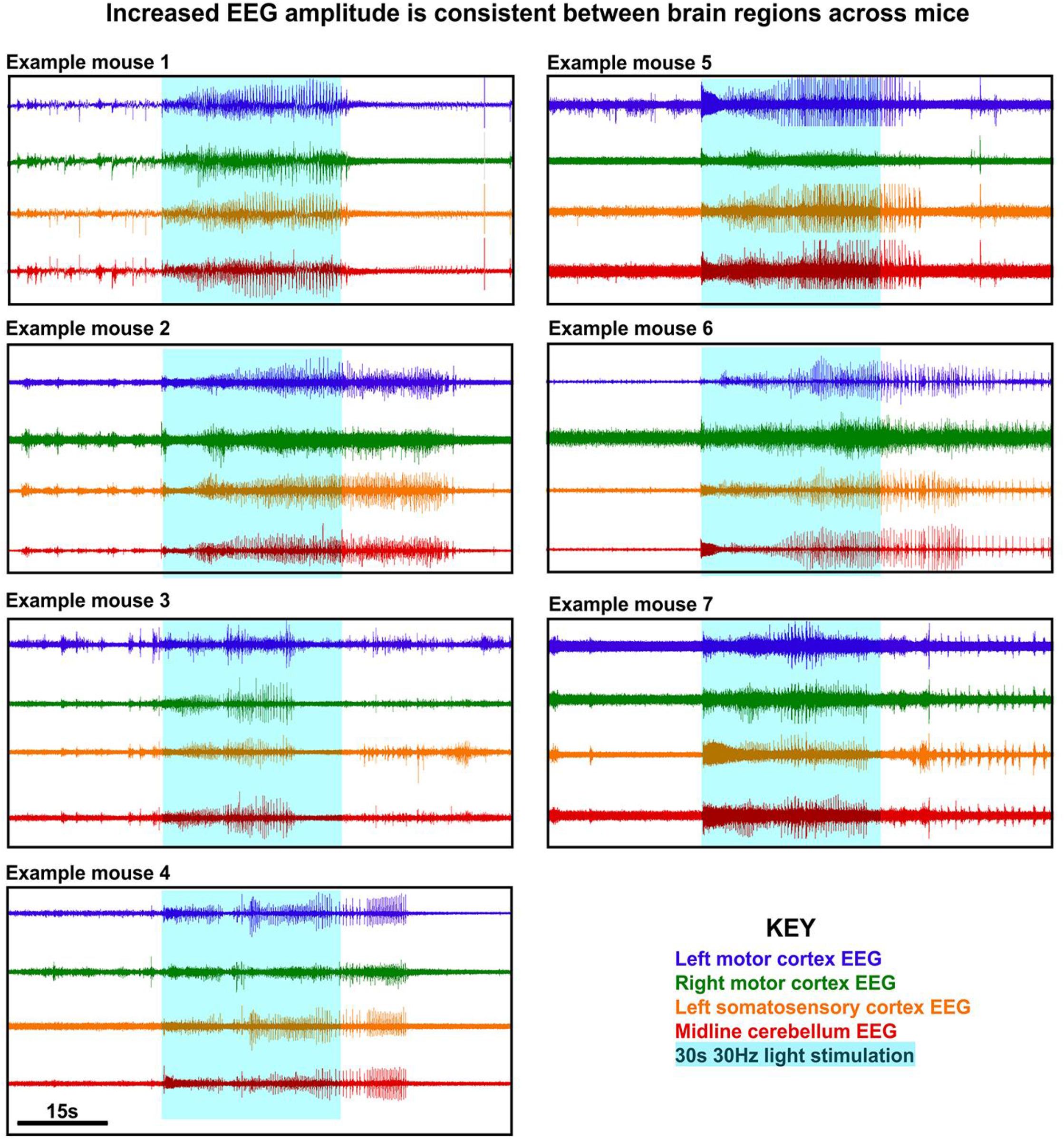
High amplitude synchrony in EEG activity is consistent across mice during optogenetically-induced seizures. Here, we show representative examples from 7 mice that all exhibited stage 6 seizures during optogenetic stimulation (blue overlayed box). A hallmark electrophysiological signature of seizure activity, high amplitude synchrony, is consistently exhibited in the left motor cortex (blue), right motor cortex (green), left somatosensory cortex (orange), and cerebellum (red).

**S6:**
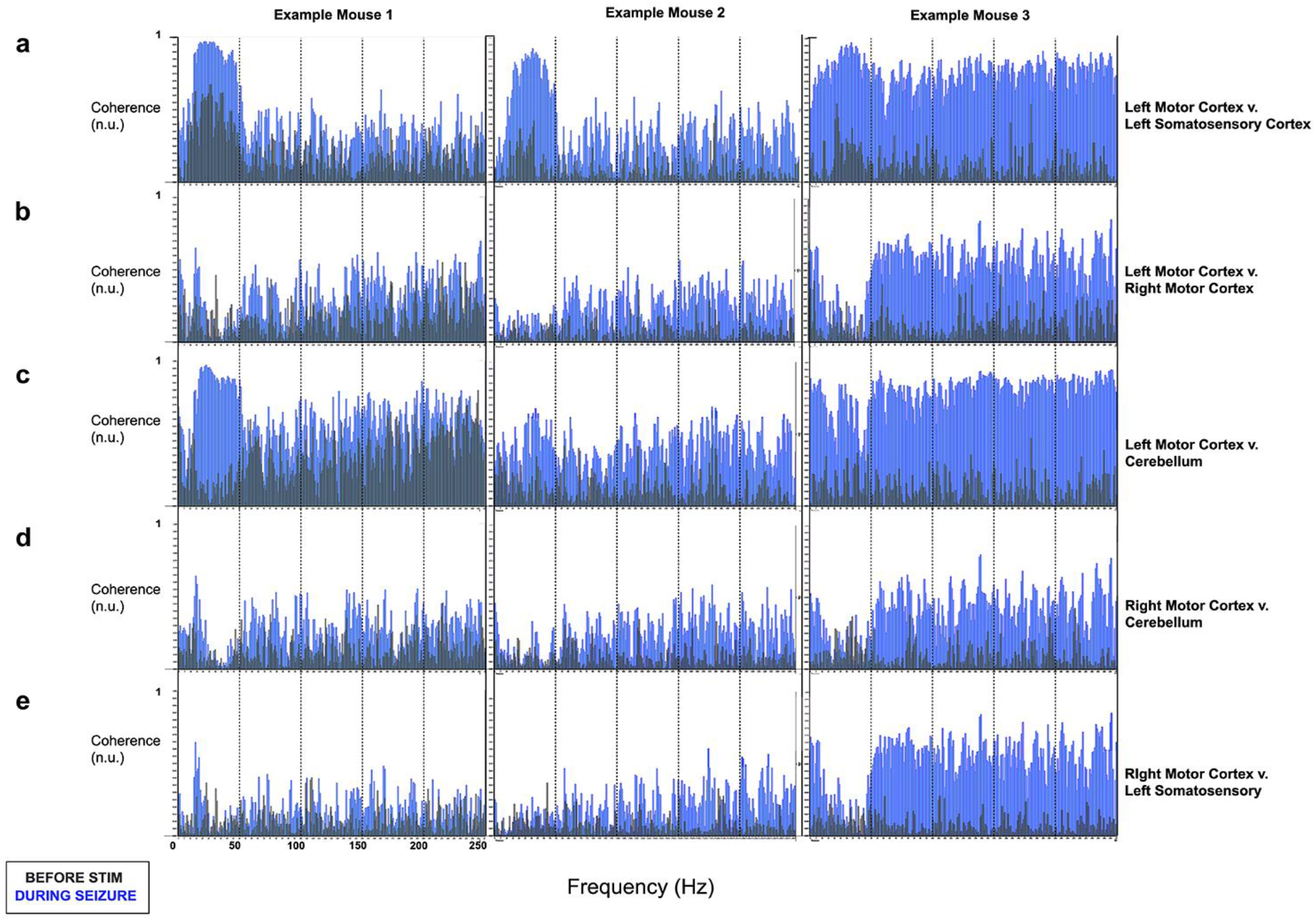
Coherence between EEG activity of left motor and left somatosensory cortex and cerebellum are most consistently increased during seizure activity. Examples of EEG coherence between brain regions from three representative mice. Here, a consistent pattern of increased coherence is seen between the before seizure (gray) and during seizure (blue) periods. (a) A large increase in coherence between left motor and left somatosensory cortex is particularly evident between 0 and 50 Hz. (b) Although coherence does increase between the right motor cortex (contralateral to the original stimulation) and the rest of the brain (d, e), this change is marginal compared to that seen in (a) and (c). (c) The left motor cortex and central cerebellum exhibit similar increases in coherence between 0 and 50Hz as that between left somatosensory and motor cortices.

**S7:**
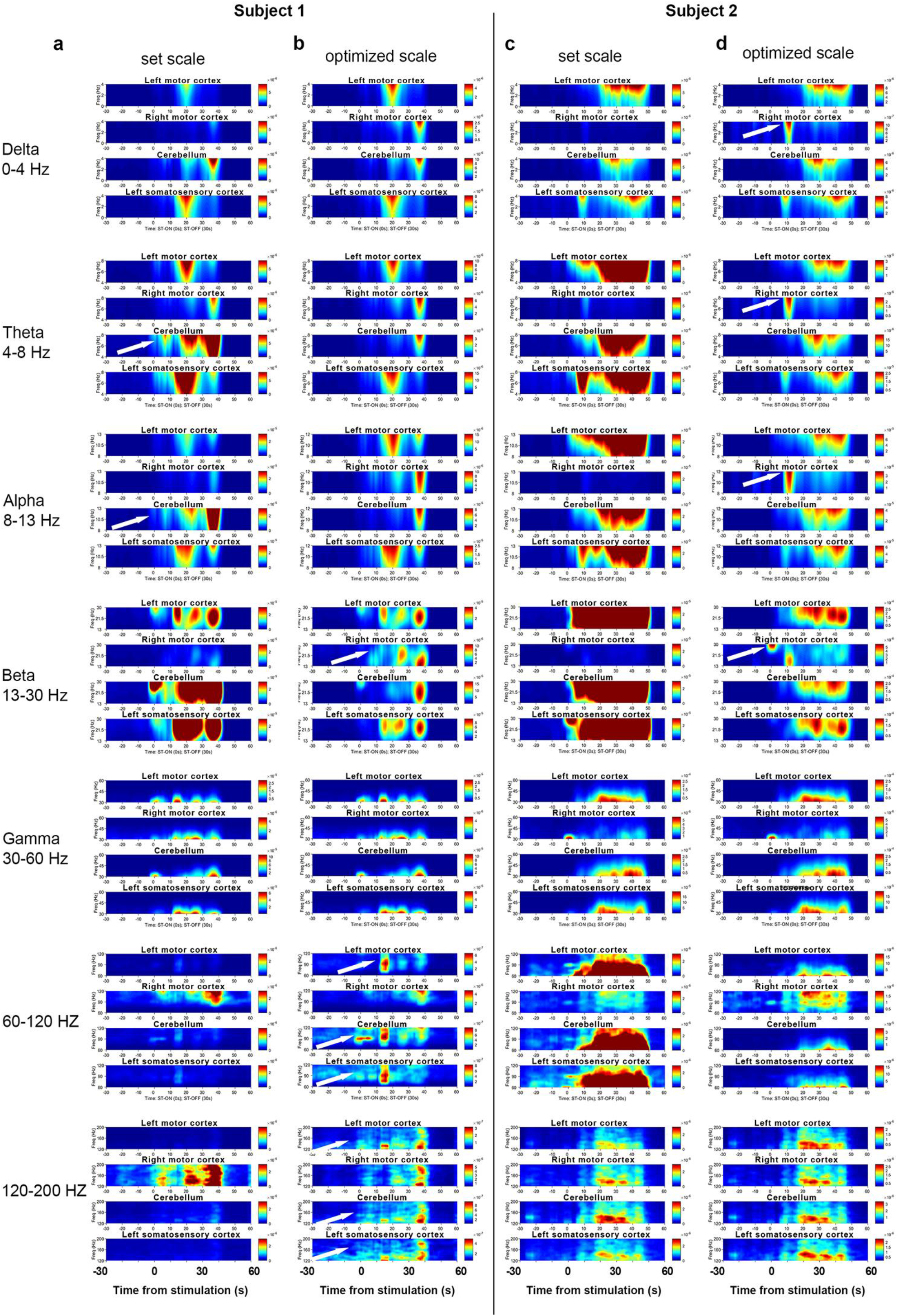
Power spectra with optimized scaling in different brain regions reveals patterned shift in activity. (a, c) Power-frequency plots from two mice experiencing seizures during the stimulation period (t = 0-30s). Due to the limitations of a set scale across brain regions, changes in activity during this period are largely obscured in several regions in Subjects 1 and 2. Here, we compare a hard/set color scale versus an optimized scale where every region receives its own color scaling according to its maximum power within the plotted frequency range. Each scaling approach reveals unique changes in activity. We show specific examples of heat plots from EEG frequencies in the delta, theta, alpha, beta, gamma, 60-120Hz, and 120-200Hz ranges. White arrows point to power changes that were previously masked by the set or optimized scaling.

**S8:**
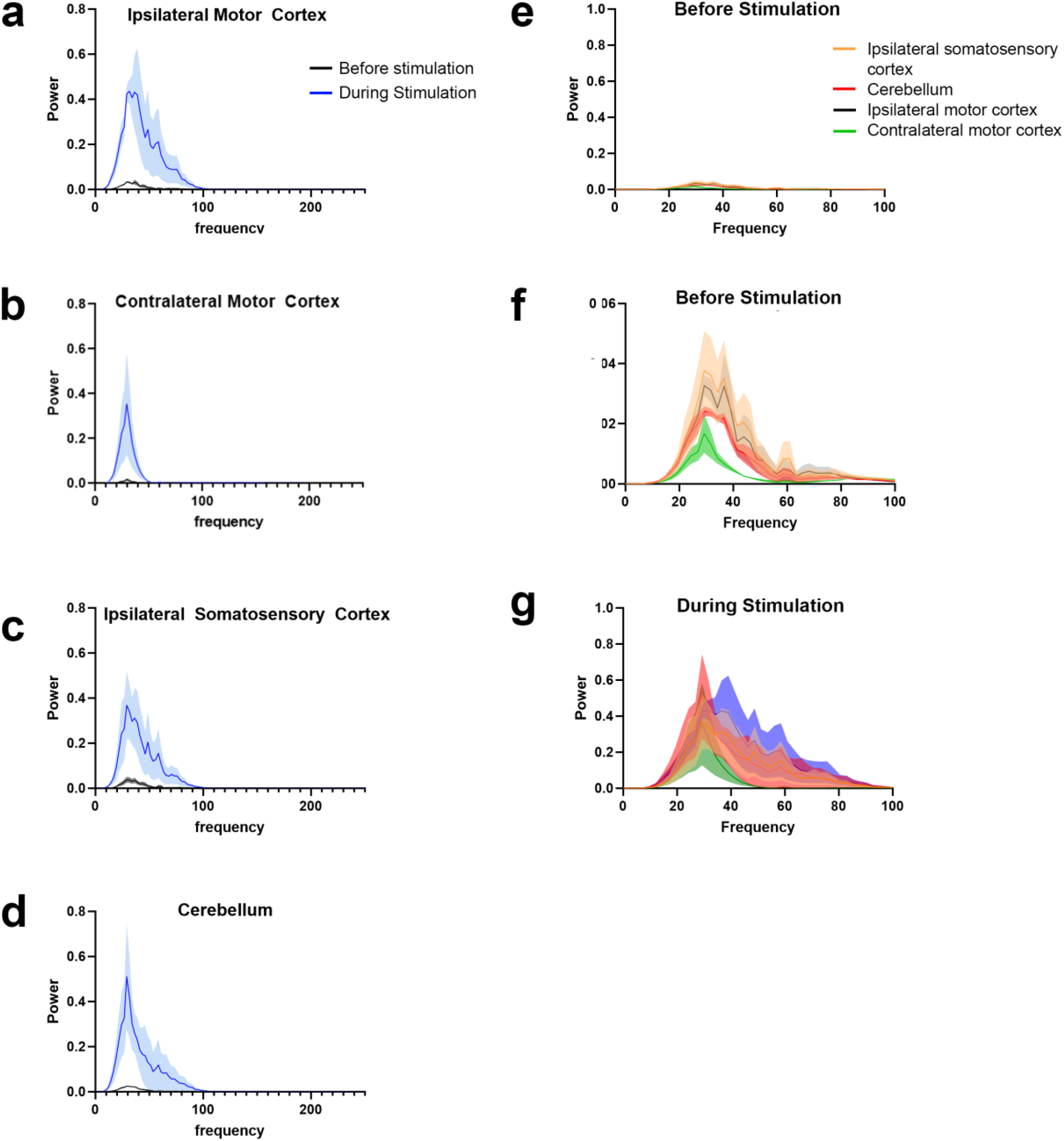
Power of frequencies under 100Hz is significantly increased during seizures. (a-d) Power plots of EcoG frequencies from 0-250Hz is shown for all four regions measured. In each region, there is a dramatic increase in the power of all frequencies between 0 and 100Hz. (e-g) Compilation of all areas onto one plot shows a comparison of changes in each brain region, demonstrating that this shift is most dramatic in the ipsilateral motor/somatosensory cortices and cerebellum, as expected due to the proximity to the site of seizure origin (left VPM).

*Video1: Optogenetic delivery of 30Hz light to the VPM elicits tonic-clonic seizures.* Video representation of a stage 6 seizure showing examples of the progression from facial and forelimb clonus to loss of posture in a mouse exhibiting a light-induced seizure. All mice, including those with severe phenotypes, experience a full recovery of general behavior after the seizures resolve.

*Video2: Tonic-clonic seizures that are induced with optogenetics are consistent and robust.* Examples of multiple mice experiencing phenotypically similar TC-seizures as a result of the unilateral, VPM-targeted 30Hz light stimulation paradigm.

*Video3: Unilateral optogenetic stimulation of the VPM is sufficient to elicit the full seizure phenotype*. Neither bilateral VPM nor co-stimulation of the VPM and cerebellar nuclei, two heavy *Cre*-expressing regions, amplifies the seizure phenotype. Therefore, to simplify surgeries and avoid unnecessary stimulation, we use unilateral light delivery to induce the seizures. Here we show an example of co-stimulation of the VPM and the contralateral interposed cerebellar nuclei.

*Video4: Optogenetic seizure induction is specific to the thalamic VPM*. Stimulation of surrounding thalamic nuclei, all *Cre-*expressing, does not elicit any overt behavioral phenotype. In this example, we show the effects of 30-Hz light delivery to the ventral posterolateral nucleus, which receives some common input with the VPM and is directly adjacent to the seizure-inducing region.

*Video5: Behavioral similarities between kainic acid-induced and optogenetic-induced seizures.* The behavioral phenotype of kainic-acid induced seizures and the optogenetic model are identical. Both exhibit forelimb and facial clonus, straub tail, and loss of body posture.

*Video6: Abnormal EEG activity is correlated with behavioral seizure.* High amplitude EEG activity that synchronizes across brain regions becomes evident as the behavioral seizure begins. For those seizures that outlast the period of light delivery, the EEG becomes increasingly erratic. When the mice recover, the EEG amplitude returns to baseline values.

*Video7: Pharmacological ablation of the cerebellar nuclei eliminates the ability to induce seizures.* Before administration of lidocaine, the mouse exhibits a severe TC-seizure upon light delivery to the VPM. After bilateral lidocaine delivery to the interposed cerebellar nuclei, light stimulation does not evoke any overt behavioral changes. However, after lidocaine washout, the seizure can once again can be elicited by light stimulation directed into the VPM.

*Table 1: Quantification of fluorescence among thalamic nuclei demonstrates regional differences in channelrhodopsin-expressing input.* Despite differences in fluorescent intensity between different thalamic nuclei such as the PO, CL, and RT, other nuclei such as the VPL do not demonstrate significant differences in levels of fluorescence when compared to the VPM. The differences in ChR2 expression alone, therefore, likely do not account for the lack of seizure behavior upon targeting of the different thalamic nuclei with fiber optics.

*Table 2: Modified Racine Scale is used to categorize and rank seizure behavior.* Using a modified Racine Scale, we compared the severity of seizure phenotypes by ranking different categorical behavioral attributes of the mice upon seizure induction with optogenetic stimulation.

