## Supplementary figures and images for "The cerebellum contributes to tonic-clonic seizures by altering neuronal activity in the ventral posteromedial nucleus (VPM) of the thalamus"

### Supplemental Figure 1

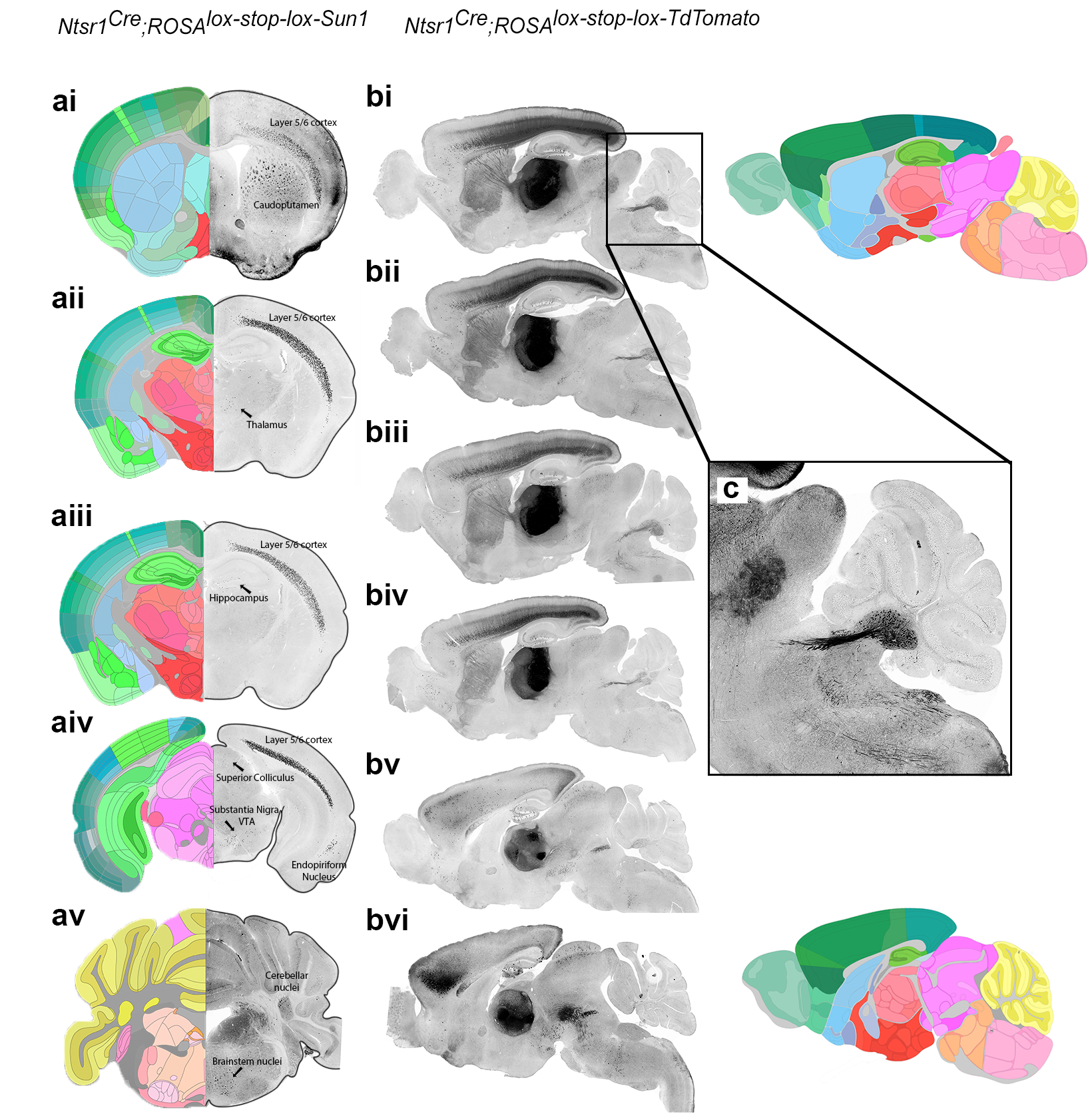

### Supplemental Figure 2

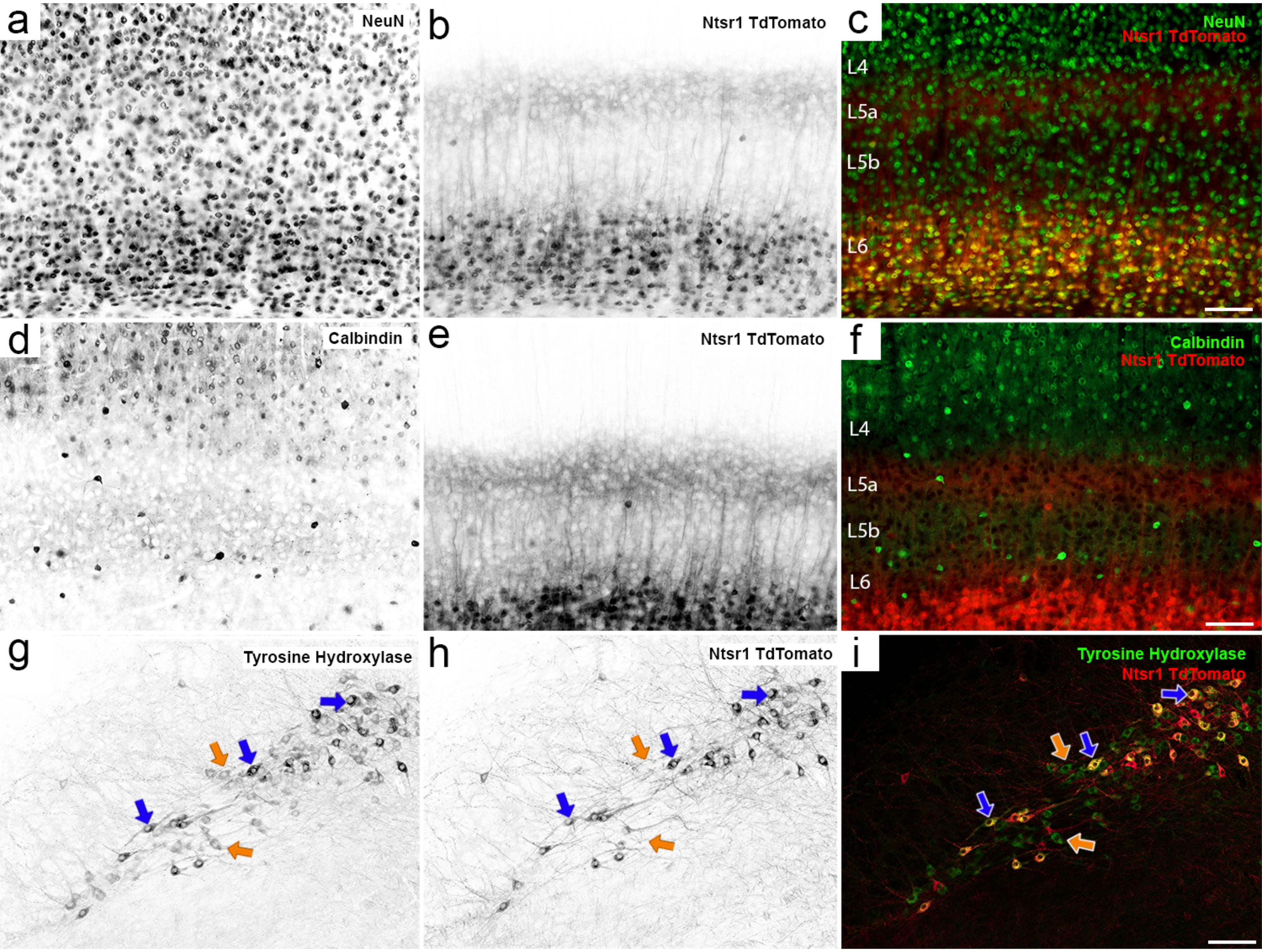

### Supplemental Figure 3

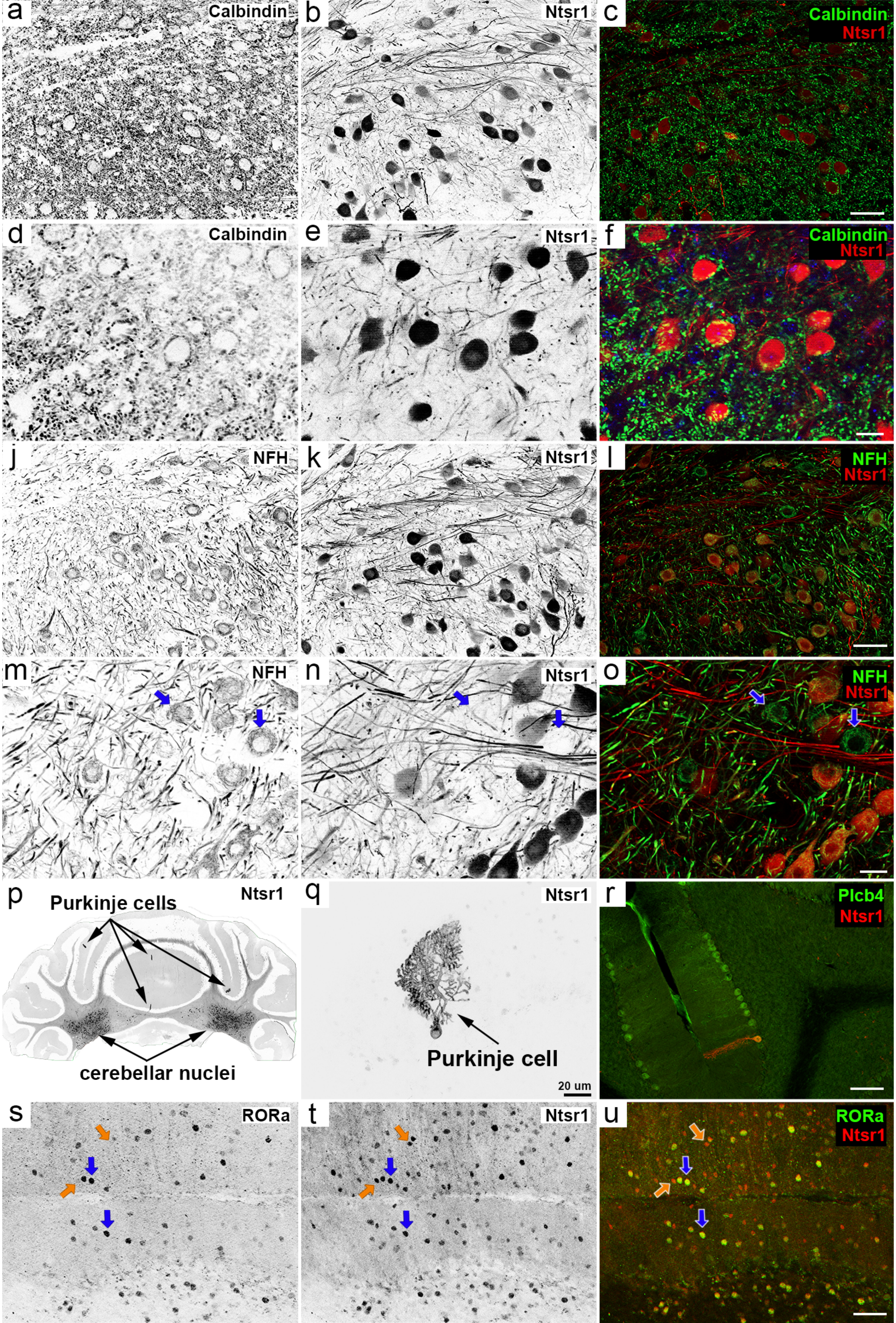

### Supplemental Figure 4

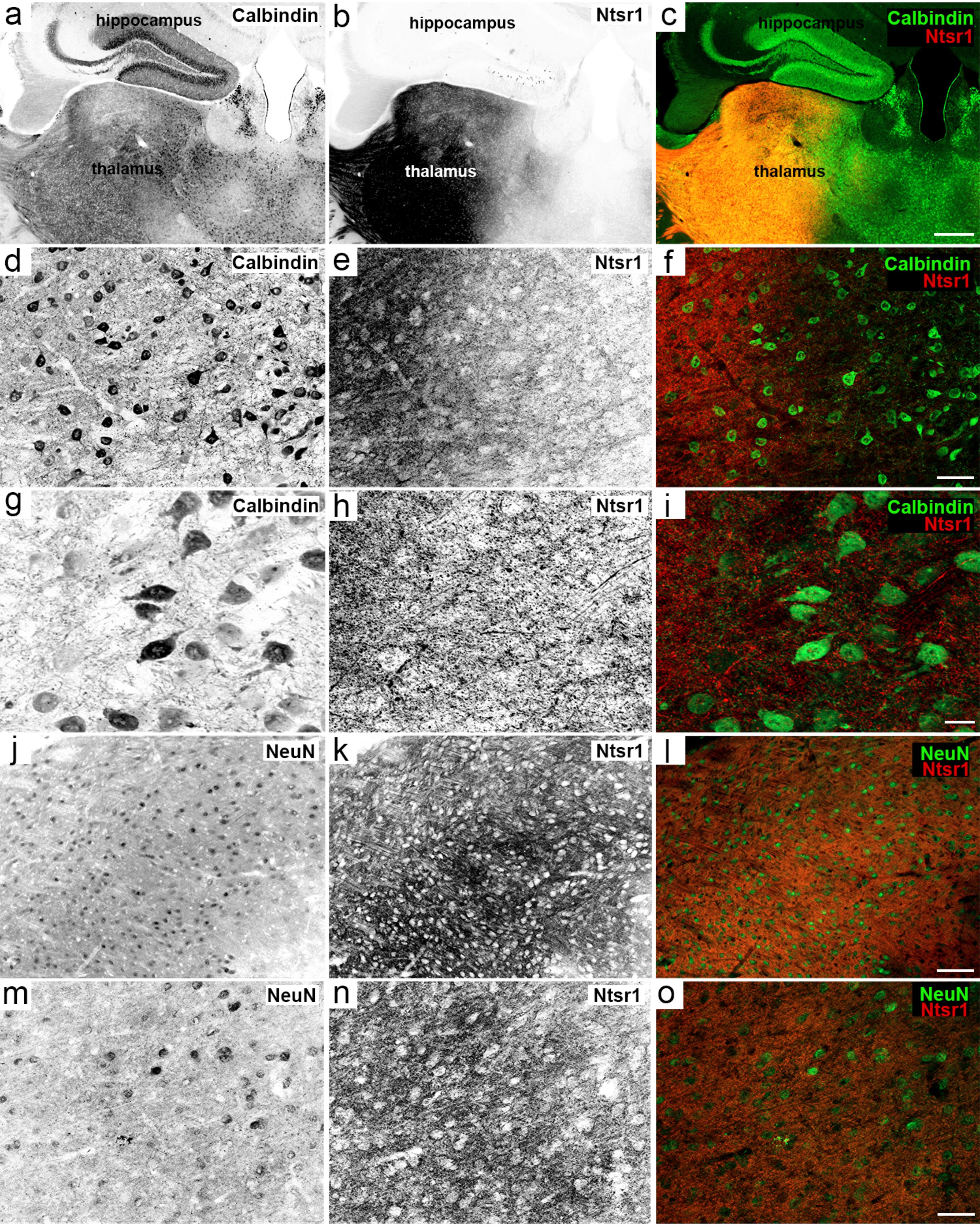

### Supplemental Figure 5

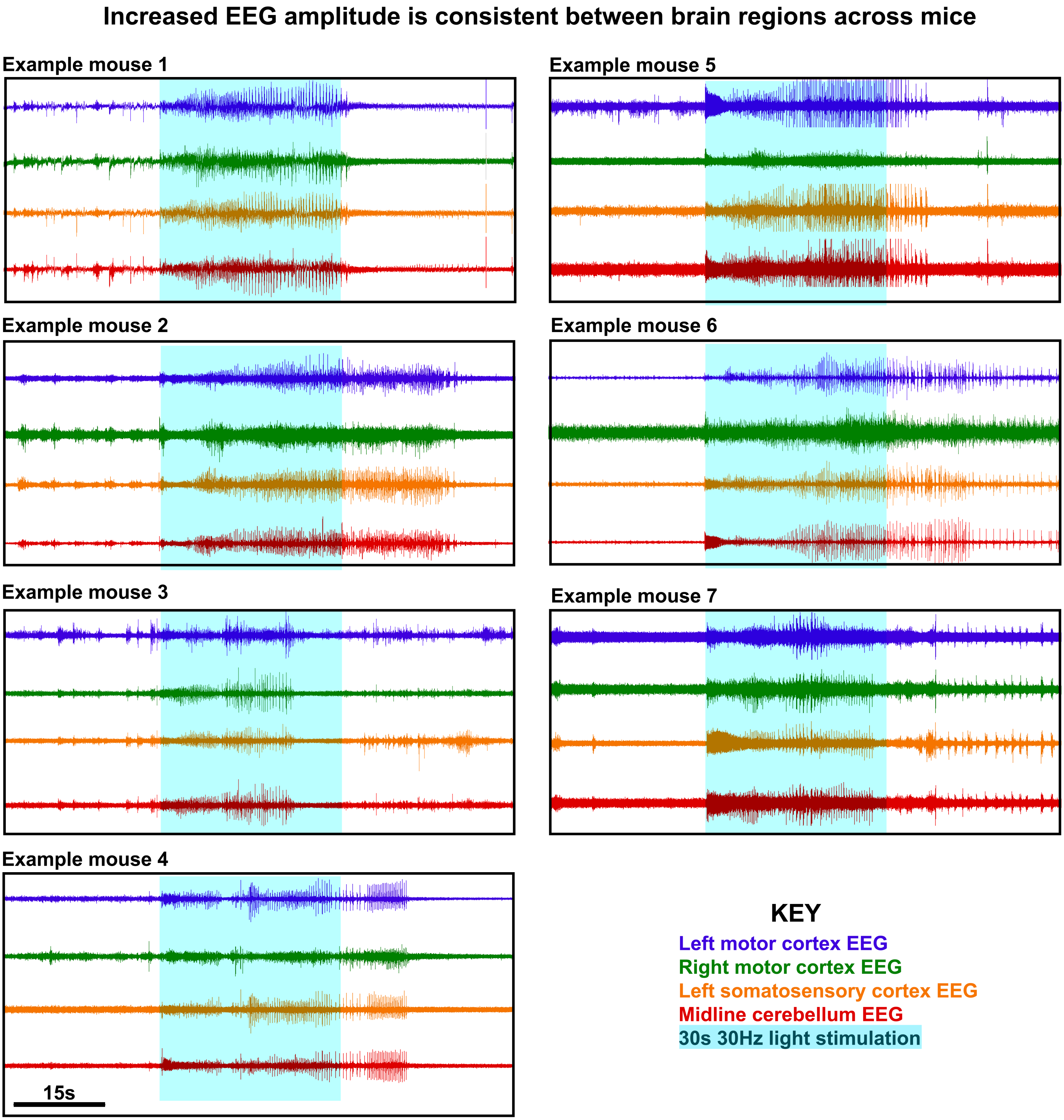

### Supplemental Figure 6

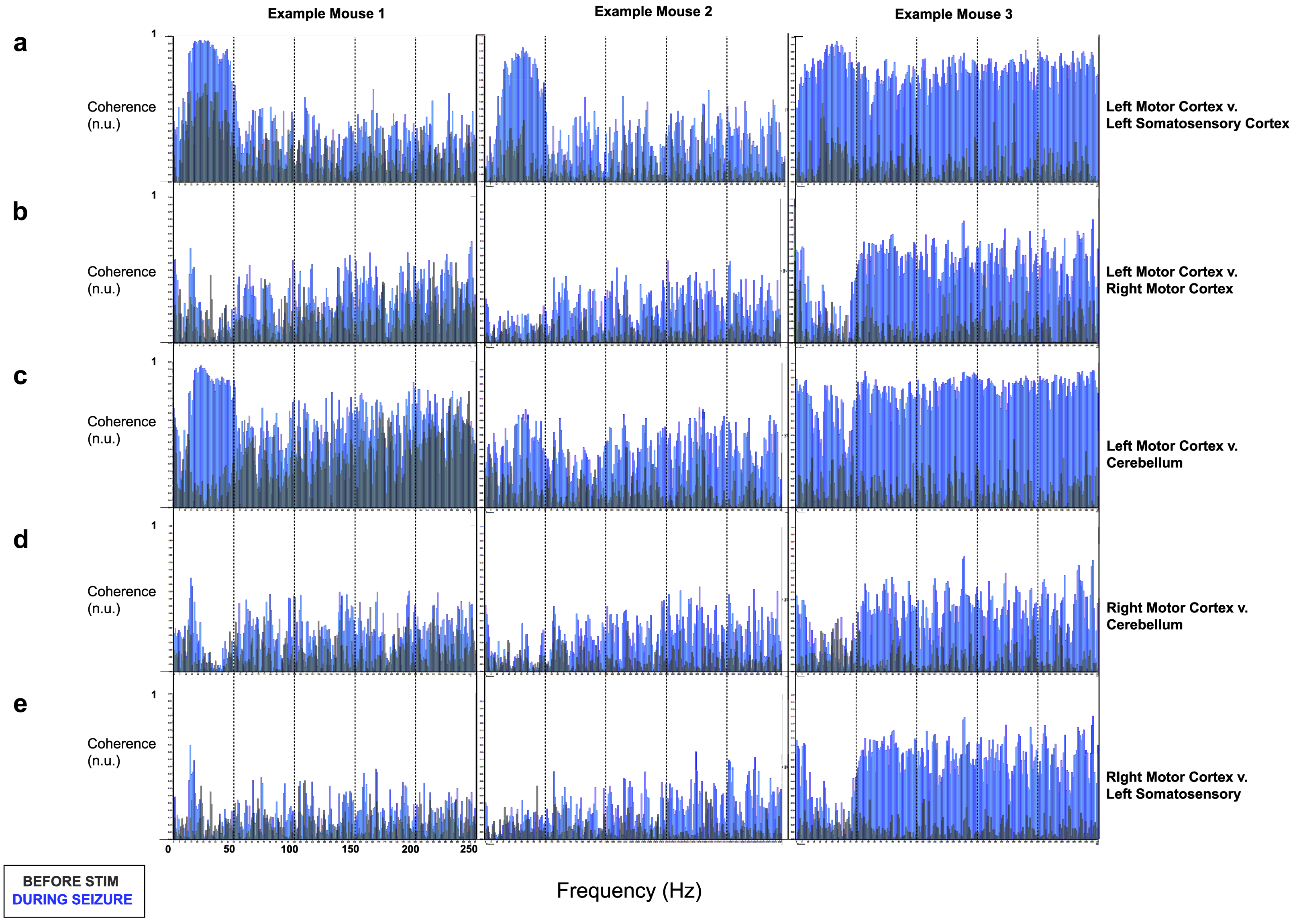

### Supplemental Figure 7

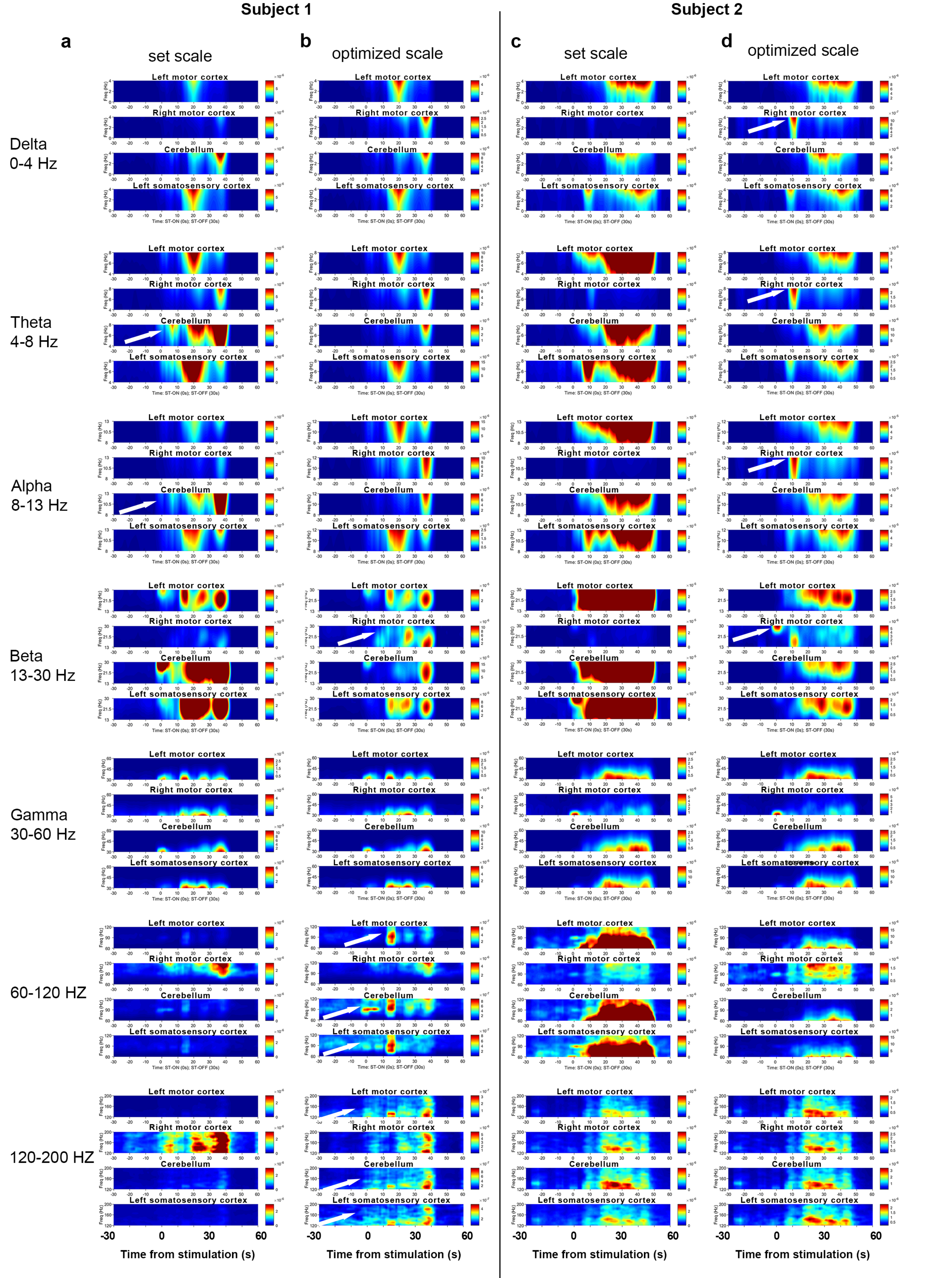

### Supplemental Figure 8

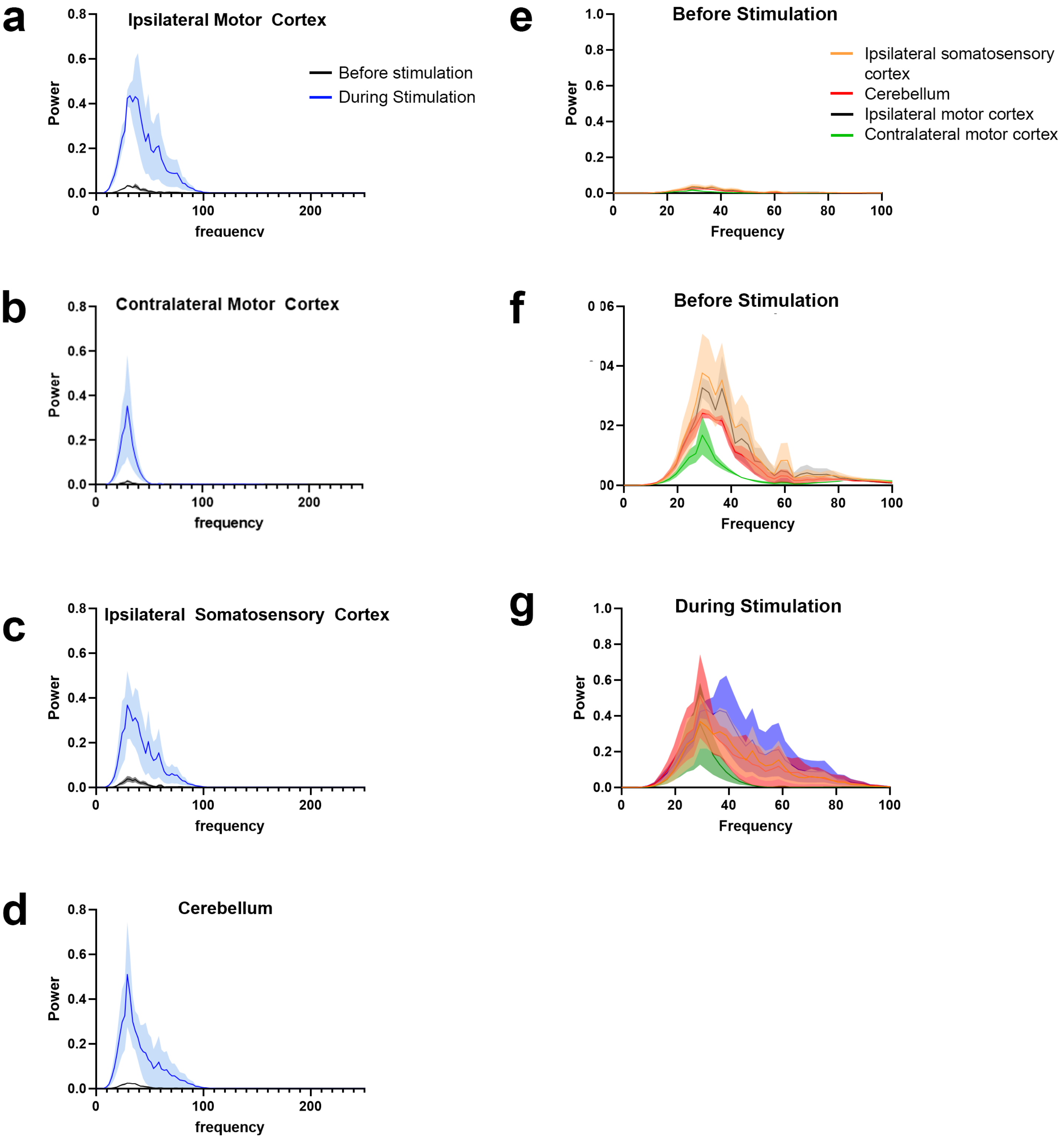
