## Supplementary material for "The cerebellum contributes to tonic-clonic seizures by altering neuronal activity in the ventral posteromedial nucleus (VPM) of the thalamus": Table 1

| Dunnett's multiple comparisons test | Mean Diff. | 95.00% CI of diff. | Below threshold? | Summary | Adjusted P Value |
|---|---|---|---|---|---|
| VPM vs. PO | 70.66 | 44.48 to 96.83 | Yes | **** | <0.0001 |
| VPM vs. LD | 77.25 | 50.71 to 103.8 | Yes | **** | <0.0001 |
| VPM vs. LP | 87.91 | 58.91 to 116.9 | Yes | **** | <0.0001 |
| VPM vs. CL | 103.8 | 76.90 to 130.8 | Yes | **** | <0.0001 |
| VPM vs. VM | 106.8 | 79.43 to 134.1 | Yes | **** | <0.0001 |
| VPM vs. VPL | -2.406 | -28.58 to 23.77 | No | ns | 0.9997 |
| VPM vs. RT | 28.81 | 2.280 to 55.35 | Yes | * | 0.0257 |
| VPM vs. VAL | 101.9 | 74.10 to 129.8 | Yes | **** | <0.0001 |
| VPM vs. LGd | -34.62 | -84.51 to 15.26 | No | ns | 0.3318 |
