## Supplementary material for "The cerebellum contributes to tonic-clonic seizures by altering neuronal activity in the ventral posteromedial nucleus (VPM) of the thalamus": Table 2

| Racine Scale | Behavioral Stage |
| --- | --- |
| 0 | no change |
| 1 | behavioral arrest, motionless staring |
| 2 | head nodding/facial clonus |
| 3 | forelimb clonus with lordotic posture |
| 4 | forlimb clonus with rearing |
| 5 | forllimb clonus with falling |
| 6 | generalized tonic-clonic activity with loss of postureal tone, wild jumping |
